# High Throughput Screening Identifies Small Molecules that Synergize with MRTX1133 Against Acquired Resistant KRAS ^G12D^ Mutated CRC

**DOI:** 10.1101/2025.06.27.662039

**Authors:** Natalie Thielen, Ning Wei, Emiko Nagai, Edward Chu, Seiya Kitamura, Chaoyuan Kuang

## Abstract

Colorectal cancer (CRC) remains a significant clinical challenge, with a 5-year survival rate of 10%. Over half of all CRCs harbor mutations in the KRAS gene, leading to poor response to standard therapy. This underscores the crucial need for novel therapeutics targeting KRAS and overcoming the growing barrier of resistance. To address these critical challenges, we conducted a high-throughput screen to identify small molecules that synergize with KRAS ^G12D^ inhibitor MRTX1133 against CRC. Through screening a 2,652 kinase inhibitor library, we discovered that Osimertinib and its analogs strongly synergize with MRTX1133 against both parental and MRTX1133-resistant cells. The top compound from the screen, NT-1, is a chemical analog of Osimertinib. NT-1 strongly synergized with MRTX1133 to suppress EGFR/MAPK signaling and induce apoptosis in an MRTX1133-resistant patient-derived organoid model of CRC. We present novel small molecule combinations with the potential to overcome the limitations of MRTX1133 with direct clinical translational applications.

**One Sentence Summary:** High throughput screening and validation in colon cancer PDOs identifies novel KRAS inhibitor combinations with potential for clinical translation.

## INTRODUCTION

Colorectal cancer (CRC) is the third deadliest cancer globally ^1^. Death due to CRC is almost exclusively caused by the progression of metastatic CRC (mCRC). The standard treatment options for mCRC are combination cytotoxic chemotherapies and biologic agents such as EGFR- and VEGF-inhibiting antibodies ^2^. However, the vast majority of mCRC patients progress through all approved treatments and eventually die from their disease. Thus, novel therapies for mCRC are desperately needed. KRAS-mutant CRC represents approximately 50% of the mCRC population. RAS oncogenes drive the activation of the mitogen-activated protein kinase (MAPK) pathway and are a principal cancer driver pathway in KRAS-mutant CRCs ^3^. These oncogenes also represent a viable treatment target, as demonstrated by numerous recent advances in KRAS-directed treatments. The KRAS **^G12D^** mutation is the most common KRAS mutation found in CRC patients, making this particular allele a high-priority therapeutic target ^4^.

Previously, the KRAS protein was considered undruggable until the FDA approved sotorasib, which targets KRAS **^G12C^** mutant tumors ^5^. This was a landmark event but also highlighted drawbacks to KRAS allele-specific inhibitors. KRAS-mutant CRCs have many adaptive and long-term resistance mechanisms, making single-agent treatment with KRAS inhibitors ineffective for long-term disease control ^6,7^. Another drawback with G12C specific inhibitors is the patients in the clinical studies experienced toxicity and acquired resistance to the point specific inhibitors. The CodeBreaK100 (NCT03600883) trial assessed sotorasib in advanced KRAS **^G12C^** mutant colorectal cancer (CRC), involving 62 patients. The study showed a 9.7% objective response rate, all partial responses ^8^. Despite not meeting the benchmark response rate, once daily sotorasib demonstrated modest anti-tumor activity and manageable safety. Many patients eventually develop progressive disease (PD) due to mechanisms of resistance that remain largely unknown. The only KRAS inhibitor currently approved for CRC patients is adagrasib, which covalently binds in the inactive form of KRAS **^G12C^** in the switch II pocket, like sotorasib. In June of 2024, adagrasib, combined with the EGFR-directed antibody, cetuximab, demonstrated promising clinical activity, resulting in accelerated FDA approval for progressive KRAS **^G12C^** mutant tumors ^9^. Moreover, KRAS **^G12C^** mutations are relatively infrequent in colorectal cancers, representing only 11% of KRAS mutations in these malignancies, and are only present in 3% of colorectal cancers overall. In contrast, KRAS **^G12D^** mutation occurs in approximately 20% of CRC patients.

More recently, the KRAS **^G12D^** inhibitor MRTX1133 was reported (**Figs. 1A**). This compound selectively binds in the switch II pocket in the G12D mutant form of KRAS (**Fig. 1B**), inhibiting its activity and thereby disrupting the downstream signaling pathways that drive tumor growth and survival. The early clinical experience with KRAS inhibitors has also revealed several challenges. There is a diversity of on-target and off-target mechanisms that can confer resistance to KRAS inhibitors and support the need for the development of additional KRAS-targeting therapeutic strategies. Many such mechanisms, including alternative KRAS mutations and mutations in other MAPK proteins, were detected in response to KRAS **^G12C^** treatment in patients. Similarly, in preclinical models of pancreatic cancer, MRTX1133 treatment led to diverse resistance mechanisms including epithelial-mesenchymal transition, PI3K-AKT-mTOR signaling, and amplification of *Kras*, *Yap1*, *Myc*, *Cdk6*, and *Abcb1a/b* ^10^. Combination therapy is a promising approach to overcome these limitations and induce deeper, more durable responses in patients^11^. Herein, through agnostic screening of over 2,600 kinase inhibitors against both parental and resistant colorectal cancer cells, we identified novel small molecule combinations that potentiate the efficacy of MRTX1133 and overcome acquired drug resistance.

**Figure 1.**
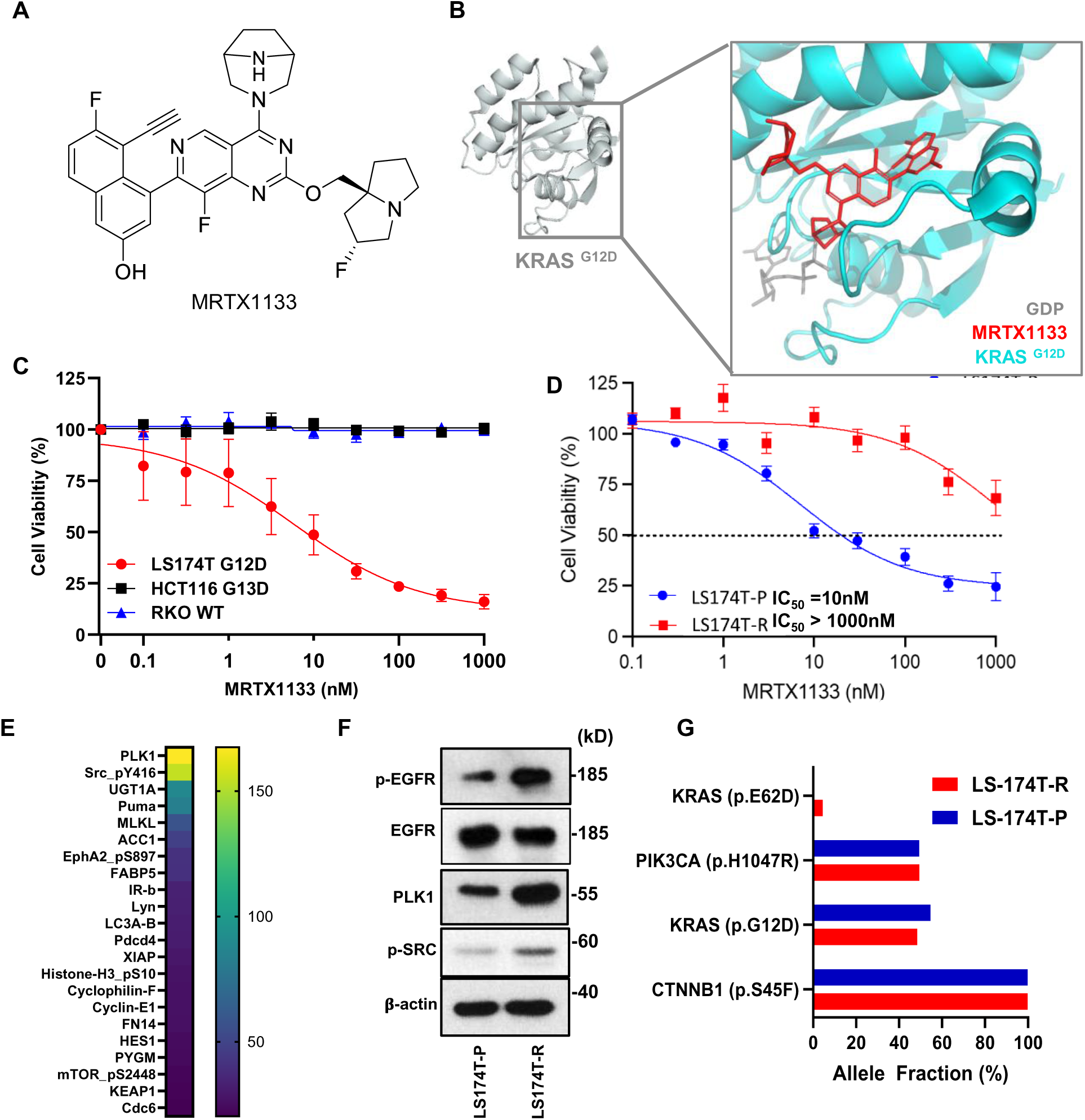
KRAS G12D inhibitor MRTX1133 and generation of resistant cell lines. **A.** Chemical structure of the KRAS G12D Inhibitor MRTX1133. **B.** KRAS ^G12D^ complex structure protein (PDB7RPZ). **C.** MRTX1133 selectively inhibits the LS-174T cells (KRAS ^G12D^ mutation) compared to HCT116 CRC cells (KRAS ^G13D^ mutation) or RKO cells (KRAS ^Wildtype^). **D.** Generation of MRTX1133-resistant LS-174T cell lines. Concentration-dependent inhibition curves of cell viability against parental (blue) or resistant (red) cells are shown. **E.** Top up-regulated gene difference between resistant and parental cell line, from RPPA data, highlighting PLK1 as most up-regulated. **F.** Immunoblot of key MAPK signaling proteins in parental and resistant LS-174T cells. **G.** Mutation profiles of parental versus resistant cells.

## RESULTS

### I. MRTX1133 resistant KRAS **^G12D^** mutant CRC cell line generation

We first validated that MRTX1133 is selectively effective against CRC cells harboring G12D mutations. Three CRC cell lines, LS-174T cells (KRAS **^G12D^**), HCT116 CRC cells (KRAS **^G13D^**), and RKO cells (KRAS **^Wildtype^**)- were treated with MRTX1133 in a concentration-dependent manner (**Fig. 1C**). As expected, MRTX1133 selectively inhibits the growth of LS-174T cells that have a KRAS **^G12D^** mutation while being ineffective against other two cell lines that do not have a G12D mutation. This is consistent with the previously reported biochemical inhibitory potency of MRTX1133 as well as previous cellular data of this compound ^12^.

To screen for drug combinations to overcome resistance, we established LS-174T cells resistant to MRTX1133. After passaging the cells eight times in the presence of MRTX1133 at 1,000 nM, we obtained cells that are resistant (R) to MRTX1133, with more than 100-fold decrease of potency compared to the parent cell line (P) (**Fig. 1D**). Resistant cell proliferation is visualized (**Supplementary Fig. S1A**), illustrating that cellular proliferation persists in the presence of up to 1 µM MRTX1133, even after 72 hours of treatment. Visual assessment of the new resistant cell line showed that it retained certain characteristics of the parental cells, such as a flat morphology when grown on tissue culture treated surfaces. (**Supplementary Fig. S1B &C**). To understand the mechanisms of resistance, we used reverse-phase protein arrays (RPPAs) and calculated the difference of protein expression between the resistant and parental cell line (**Fig. 1E**). The antibody repertoire covers key oncogenic pathways such as PI3K/AKT, RAS/MAPK, Src/FAK, TGF-β/SMAD, JAK/ STAT, DNA damage repair, Hippo, cell cycle, apoptosis, histone modification, and immune oncology. Analysis of the RPPA data revealed significant upregulation of protein levels, notably PLK1 and Src in the resistant LS-174T cell line. Conversely, there were a number of proteins, including tumor suppressors, that were down-regulated in the resistance cell line, indicating multiple potential mechanisms of resistance at the protein level (**Supplementary Fig. S1D**). For upregulated proteins identified in RPPA, we performed an immunoblotting analysis to confirm the increased protein levels in the resistant cells compared to the parental cells (**Fig. 1F**). In addition to proteins identified from RPPA analysis, the resistant cell line exhibited upregulation of phospho-EGFR (p-Y1068) expression, aligning with previous studies that confirmed the EGFR feedback mechanism upon treatment with MRTX1133 ^13^. Furthermore, genomic mutational profiling identified the emergence of a new KRAS mutation in the resistant cell line (**Fig. 1G**), absent in the parental line. This newly acquired mutation may, at least in part, contribute to the observed resistance phenotype and alterations in signaling dynamics within the resistant LS-174T cells. These results indicate that prolonged treatment with MRTX1133 can lead to diverse genomic and proteomic resistance mechanisms in CRC. Our pair of parental and resistant cell lines also serve as useful tools for discovery and validation of therapies to overcome resistance, as follows.

### II. High-throughput small molecule screening and triage

We conducted a high-throughput screen to identify compounds that (i) potentiate MRTX1133 efficacy in colorectal cancer cells, and (ii) overcome resistance. We hypothesized that the inhibition of kinases in the cancer cell signaling pathway potentiates the efficacy of the KRAS **^G12D^**inhibitor and helps overcome resistant mechanisms. A kinase inhibitor library consisting of 2,652 compounds was selected for our screening campaign. The library contains inhibitors targeting various protein kinases and lipid kinases. Detailed information of the library including the compound structures and target protein/pathways is proved in the supplementary information (**Supplementary Table S1**) To identify compounds that are effective against resistant cells and those that synergize with MRTX1133, screening was performed under four distinct biological conditions; parental *vs* resistant LS-174T cells in the presence or absence of MRTX1133 (**Fig. 2A**). The compound library was dispensed at a final concentration of 10 nM onto cells seeded in 384-well plates and cultured for 72 hours (**Supplementary Fig. S2A**). The concentration of MRTX1133 in the screening was carefully selected based on its IC_50_ values in both parental and resistant cell lines (**Fig. 1D**), ensuring that its effect remained below the inhibition cutoff in the primary screen while still allowing the identification of potential synergistic compounds.

**Figure 2.**
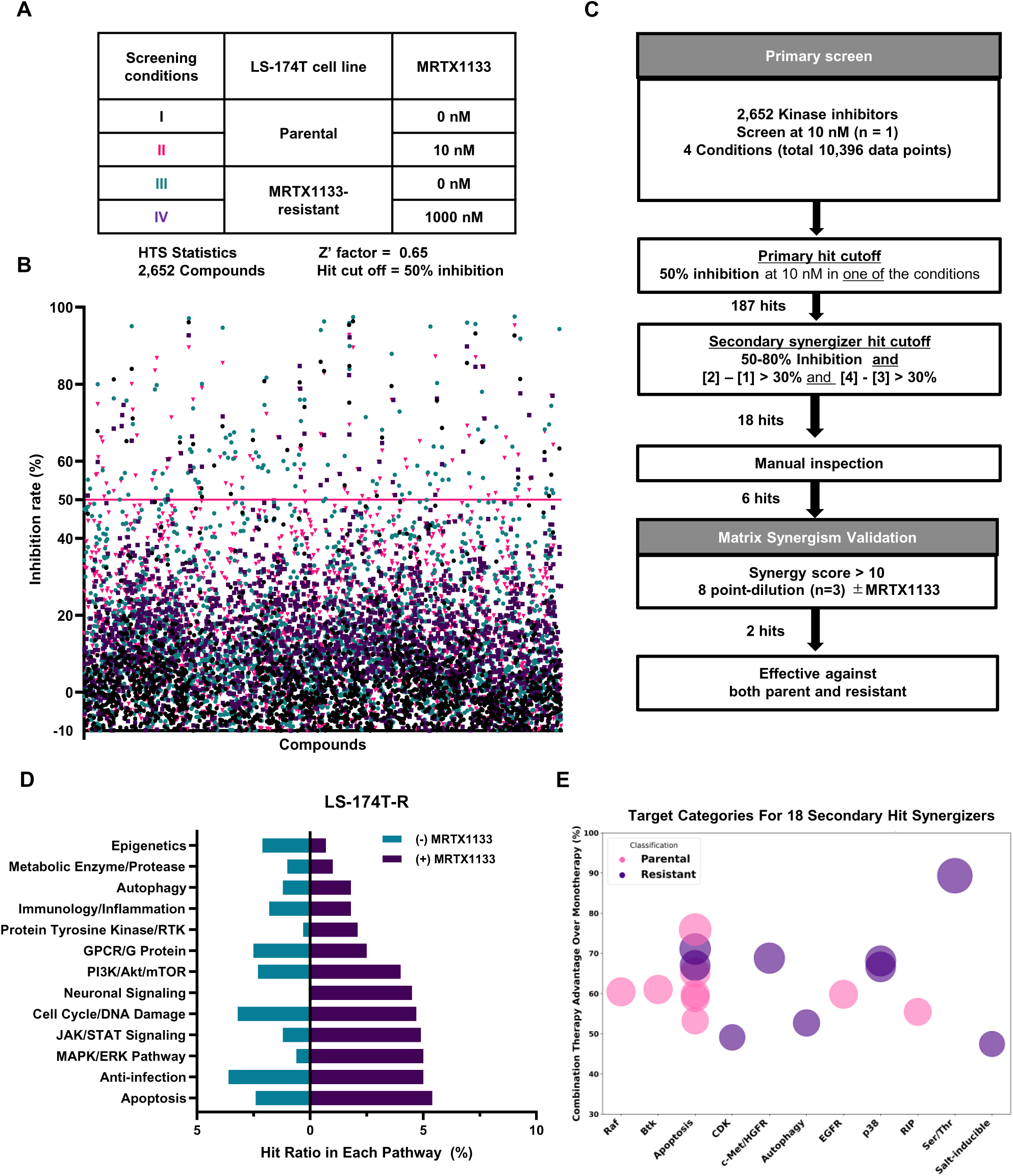
Small molecule screening against LS-174T Parental (P) and MRTX1133-resistant (R) in the presence or absence of MRTX1133. **A.** Primary screening conditions. A total of four conditions were tested with 2,652 small molecule kinase inhibitor compounds at 10nM, leading to 10,608 data points. **B.** Scatter plot of LS-174T parental (black: -MRTX1133, red: +MRTX1133) and resistant (green: -MRTX1133, purple: +MRTX1133) cell lines. The cell viability was measured after 72 hours in the presence or absence of MRTX1133 at 10nM (parental) or at 1,000nM (resistant). The inhibition rate was calculated based on the vehicle treatment control in each group. Symbol colors match screening conditions stated in 2A. **C.** Hit triage to prioritize synergizers in the screen. **D.** The hit ratio in each biological pathway in LS-174T-R cells. Ratios calculated as number of hits normalized to the percent representation of that pathway within the library. **E.** Eighteen secondary hits (see 2C) are graphed based on the advantage of combination treatments (II and IV) over monotherapy treatment (I and III).

The primary screening scatter plot is shown in **Fig. 2B**. The screening assay performed well, with an overall Z’ factor exceeding 0.6. Each group yielded distinct hit compounds, with the monotherapy-treated groups showing approximately a 1% hit rate (**Supplementary Fig. S2B**). Among the 2,652 compounds tested, 187 compounds met the primary hit cutoff of 50% inhibition in at least one condition, resulting in an overall primary hit rate of 7%. To identify synergizers of MRTX1133 and to avoid general cytotoxic compounds, secondary hit cutoff criteria were applied; compounds were selected if they exhibited at least a 30% increase in inhibition in the MRTX1133 combination groups (conditions **II** and **IV**) compared to their corresponding monotherapy groups (conditions **I** and **III**). Details of the hit categories are provided in **Supplementary Table S2**. After manually inspecting the hit molecules to exclude pan-assay interference and non-drug-like compounds and selection based on mechanistic insights and clinical relevance, six compounds were chosen for further biological evaluation of their synergistic activity.

### III. Validation of small molecule combinations for overcoming MRTX1133 resistance

Dose-response matrices were created for all six selected compounds to validate their synergistic activity with MRTX1133 (**Supplementary Fig. S3A**). Based on the cell viability data in a matrix format, we calculated the ZIP synergy score to quantitatively assess synergy between each compound and MRTX1133^14^. Three compounds showed strong synergism with MRTX1133 against both parent and resistant cells. It is interesting to note that all four validated molecules target the MAPK pathway, including mutant EGFR inhibitor, Limeritinib (EGFR inhibitor), and p38α inhibitor (Fig. 3A.). The ERKi showed synergy in the parental cell line at higher concentrations, but less synergy in the resistant cell line. The transcriptional inhibitors CDK12-IN-E9 and CDK8-IN-4 showed limited synergism, even when tested at lower concentrations matrices (**Supplementary Fig. S3B).**

**Figure 3.**
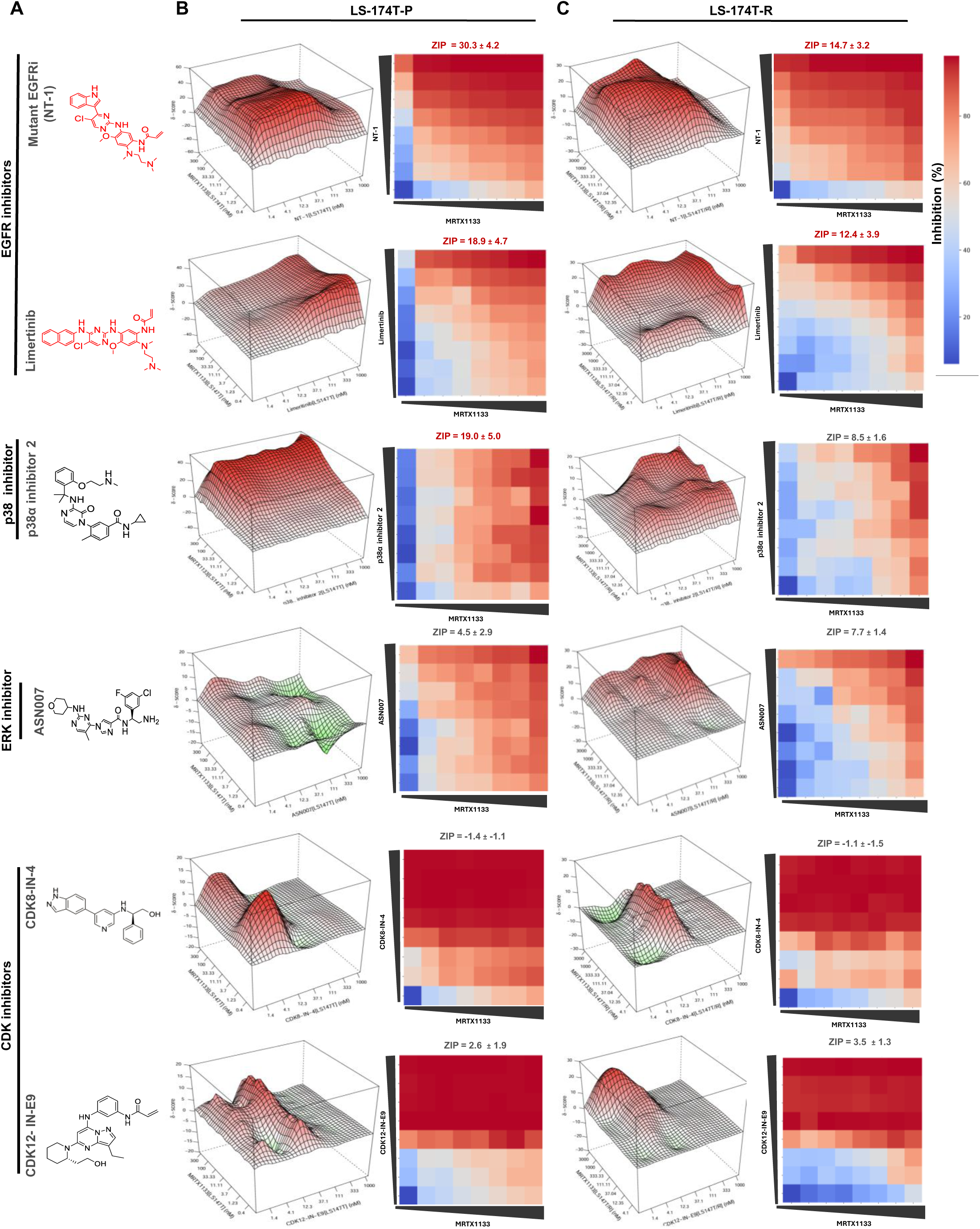
Matrix validation of the synergistic effects of top hit compounds in combination with MRTX1133. **A.** Chemical structures and names of the top compounds, with the most synergistic highlighted in red. **B.** Synergy diagram of top 6 hits treated with MRTX1133 analyzed by R package “synergyfinder3.0” (3D plots on left). LS-174T-P cells were treated for 72h with various concentrations of the indicated inhibitors. The concentrations of hits or MRTX1133 were used in a three-fold dilution series (1.4, 4.1, 12.3, 37.1, 111.1, 333.3, and 1000nM for hits; 0.4, 1.2, 3.7, 11.1, 33.3, 100, and 300nM for MRTX1133). Cell viability was measured and normalized to DMSO (control) treated cells (2D plots on right). ZIP values were simulated using zero interaction potency model analysis. **C.** LS-174T-R cells were treated for 72h with various concentrations of the indicated inhibitors. Synergy diagram of top 6 hits in LS174T-R cells were treated for 72h with same concentration of the hits with MRTX1133 used in a three-fold dilution series (12.3, 37.1, 111.1, 333.3, 1000, and 3000nM). Same analysis as 3B.

Both parental and resistant cell lines revealed robust synergy between the irreversible EGFR inhibitors and MRTX1133, which was further supported by the 3D synergy matrices. Notably, two compounds—Mutant EGFRi and Limeritinib—demonstrated synergy scores above 10, signifying strong synergistic potential, highlighted in red (**Fig. 3B** & C). Mutant EGFRi, an experimental inhibitor with no formal name ^15^ will be referred to as “NT-1” throughout the study to ensure clarity. NT-1 exhibited potent synergistic effects in both the parental and resistant colorectal cancer (CRC) models, positioning it as a promising candidate for further exploration. Additionally, while the p38α and ERK inhibitor combinations had increased ZIP synergy scores, they did not achieve scores in the synergistic range of greater than 10. These inhibitors’ potential therapeutic value remains a topic of ongoing investigation. Based on these findings, we have prioritized the EGFR inhibitors for subsequent biological validation and therapeutic development, as they offer the most promising synergy in the context of overcoming resistance to MRTX1133.

### IV. Comparison of EGFR inhibitors as MRTX1133 synergizers in vitro

Given previous literature, it is unsurprising that the top hits that emerged were small molecule EGFR inhibitors. Inhibition of mutant BRAF^V600E^ with BRAF kinase inhibitors, and inhibition of KRAS^G12C^ or ^G12D^ with allele specific inhibitors, reliably triggers the adaptive mechanism of EGFR upregulation and subsequent increase in mitogenic signaling ^16^ . This has led to the expectation that co-inhibition of EGFR and KRAS would lead to improved responses. These findings provide strong evidence for synergy, yet when we look at the complete list of EGFR inhibitors found in the kinase library (**Supplementary Fig. S4A**), there is a wide variety of inhibition, leading to some inhibitors working much better than others, suggesting a more complex interaction than initially anticipated. Interestingly, unlike in non-small cell lung cancer, CRC has an incredibly low rate of EGFR mutation (<0.5%) ^17^. In both the LS174T-P and -R models used in our screen, EGFR mutations were not detected (**Fig. 1H**). To investigate the hypothesis that co-inhibition of EGFR is the primary mechanism of synergy, we assessed the interaction between MRTX1133 and a panel of well-characterized EGFR inhibitors. The top hit, NT-1, was found to be structurally similar to Osimertinib (**Fig. 4A**), with the minor chemical differences shown in red. Despite its inclusion in the kinase library, Osimertinib did not emerge as a hit (**Fig. 4B**), likely due to the screening concentration and hit cutoff used. The cellular potency of NT-1^18^ was further characterized against previous literature on Osimertinib^19^, Limeritinib^20^, and Gefitinib^19^, the latter being an FDA-approved first generation wild-type EGFR inhibitor also included in the screen (**Fig. 4C**). We tested NT-1 and Osimertinib in a dose-response matrix, as shown in Figure 3, to compare their functionality across multiple KRAS **^G12D^** cell lines. It appears that Osimertinib synergized with MRTX2233 in LS-74T-P and LS-513 and almost reached the ZIP score cutoff of 10 in LS-174T-R (**Fig. 4D**). Comparing Osimertinib to its analog NT-1 (**Fig. 4E**), we observed the highest point of synergy (boxed in yellow on the 2D plot) for NT-1 is in the low nanomolar (<100 nM) range compared to Osimertinib (∼100 nM). To further investigate this finding, we compared the four (EGFR inhibitor) EGFRi’s individually and in combination with MRTX1133 across the three different KRAS **^G12D^** cell lines in a dose-response experiment for 72 hours (**Fig. 4F-H**) The EGFR inhibitors were treated beginning with 100nM decreasing by 2-fold and MRTX1133 began at 20 nM (P) or 2,000 nM (R). Comparing the combination treatment and monotherapy treatment we see a significant difference in fold change for the NT-1 combination groups compared to the other small molecule inhibitors. Limeritinib showed some significance in our testing which confirms its potency in the screen. We observed a significantly greater efficacy with NT-1, which was markedly more effective than the other inhibitors at 100nM over 72 hours. We further validated the potency and synergy of NT-1 and MRTX1133 in a decreased dose-response matrices (**Fig. 4I**). NT-1 was the only EGFRi that reached the synergistic ZIP score cut-off of 10 in both LS-174-P and LS-174-R cells Synergy was also observed between Limeritinib and MRTX1133 in LS-513 cells. Overall, NT-1 outperforms the other small molecule inhibitors as a combination therapy. To further study the selectivity and potency of NT-1, we tested biochemical kinase selectivity of NT-1 starting at 1 µM. (**Fig 4J**.) There was a range of potency by NT-1 with greater potency toward EGFR ^wildtype^ and EGFR ^T790M^. PLK-1 and Src seem like unlikely targets for synergy but were tested based on their upregulation in RPPA data. Overall, our results suggest that NT-1 is a uniquely synergistic molecule when combined with MRTX1133.

**Figure 4.**
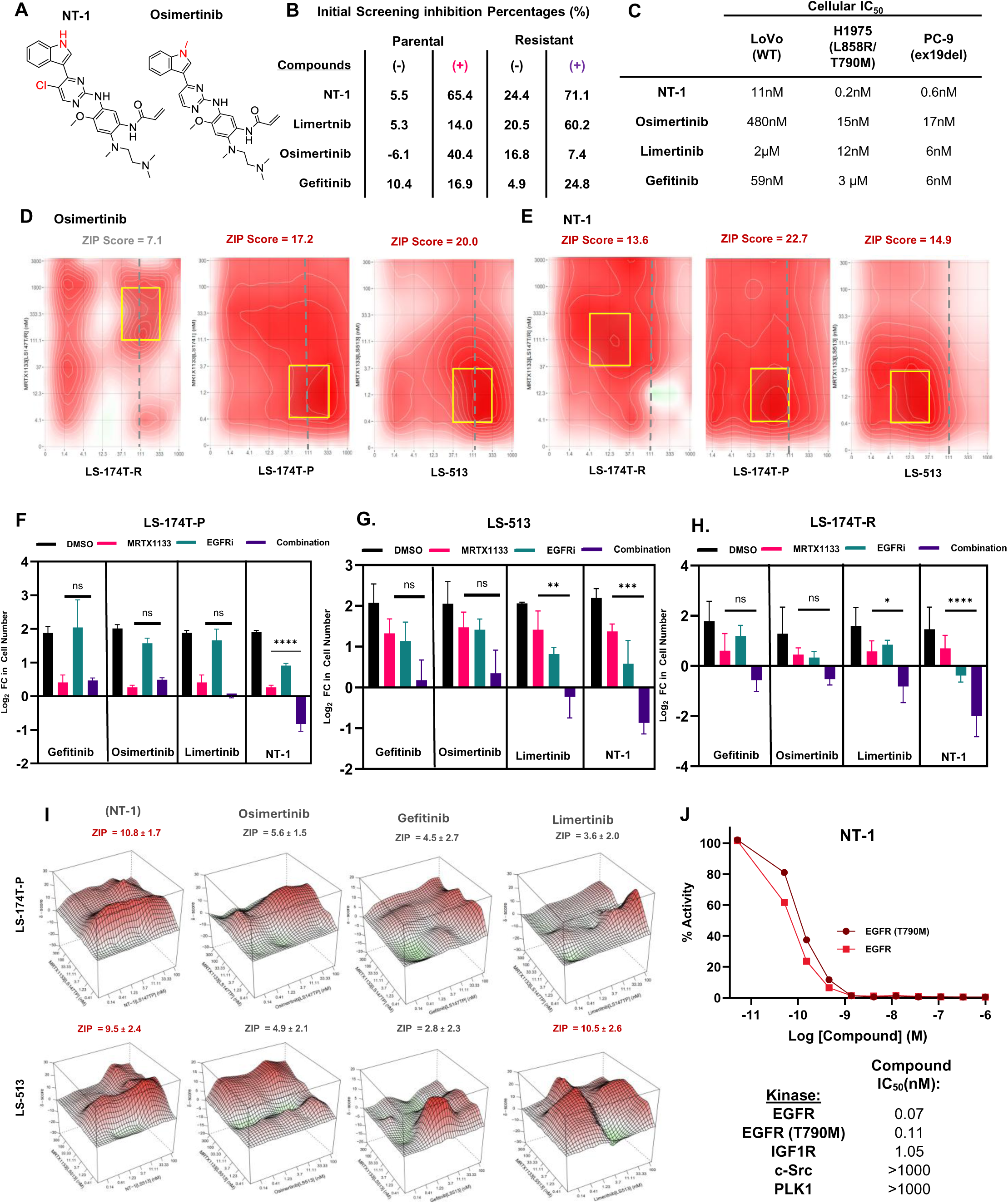
NT-1 is a potent inhibitor of EGFR and strongly synergizes with MRTX1133 to treat KRAS G12D CRC cells. **A.** The chemical structures of NT-1 and Osimertinib, differences highlighted in red. **B**. Table of top two hits initial screening percentages alongside Osimertinib and Gefitinib. **C**. Wildtype and mutant EGFR inhibitors with associated cellular IC_50_ data. **D & E**. 2D synergy plot matrices in three different KRAS ^G12D^ CRC cell lines, comparing Osimertinib and NT-1, most synergistic area highlighted in yellow. Grey dotted line marks 111nM, showing NT-1 works below 100nM in most synergistic areas. **F-H.** Proliferation assay over 3 days in a panel of three KRAS **^G12D^** CRC cell lines; LS-513 (F), LS-174T-P (G), and LS-174T-R (H) treated with DMSO, MRTX1133 (20nM (F & G) and 2000nM (H), Gefitinib(100nM), Osimertinib (100nM), Limertinib, (100nM), NT-1 (100nM) or combination. Graph reflects the relative change in cell number (log2 fold scale to best visualize loss of cells) compared with day 0. **I.** Synergy diagram of NT-1, Osimertinib, Limertinib, and Gefitinib treated with MRTX1133 analyzed by R package “synergyfinder3.0”. LS-174T-P and LS-174T-R cells were treated for 72h with various concentrations of the indicated inhibitors. The concentrations of EGFRi or MRTX1133 were used in a three-fold dilution series (3.13, 6.25, 12.5, 25, 50, and 100μg/mL for cetuximab; 3.13, 6.25, 12.5, 25, 50, and 100nM for Gefitinib, Limertinib, and NT-1. ZIP values were simulated using zero interaction potency model analysis. **J**. Biochemical data of NT-1 reveals kinase targets, showing NT-1’s highest potency potent towards EGFR ^T790M^ and EGFR ^Wildtype^.

### V. Combination Therapy Blocks Downstream MAPK Pathway in MRTX1133 Resistant Cells

We next studied the synergistic effect of NT-1 (**Fig. 5A**) compared to Cetuximab (**Fig. 5B**), an EGFR antibody that was recently approved for combination treatment with adagrasib (KRAS **^G12C^** inhibitor) for colorectal cancer ^21^. We utilized the Chou-Talalay median effect method ^22^ to determine the combination index (CI) response of NT-1 (average CI = **0.6**) and Cetuximab (average CI = **0.9**) in combination with MRTX1133. The LS-174T-R cell line showed an increased synergistic effect with NT-1 compared to Cetuximab, especially when comparing the difference in CI values (**Fig. 5C**). We next investigated the molecular mechanism underlying the strong synergy between EGFR inhibition by NT-1 and MRTX1133, focusing on the resistant cell line. The initial steps started with monotherapy of NT-1 to determine effective concentration after 24 hours (**Supplementary Fig. S5A**). We begin to see suppression of EGFR and increased apoptosis around 60 nM. The combination treatment moving forward included 60 nM of NT-1. MRTX1133 is known to downregulate the ERBB receptor feedback inhibitor 1 (ERRFI1), which is a negative regulator of EGFR, causing feedback activation. The EGFR family comprises four distinct membrane tyrosine kinase receptors: EGFR/ErbB-1, HER2/ErbB-2, HER3/ErbB-3, and HER4/ErbB-4, which are activated upon ligand binding to the extracellular domain of these receptors ^13,23,24^. This induction of ERBB2 and ERBB3 expression, in turn, reactivates KRAS signaling, limiting the sensitivity to these single-agent drug treatments. The monotherapy of MRTX1133 and the combined effect of NT-1 and MRTX1133 in the resistant cells are shown in **Figure 5D**. There is complete suppression of p-EGFR in the combination groups, as well as downstream suppression of p-ERK and an increase in Caspase 3, indicating that the cells are undergoing apoptosis with this combination treatment.

**Figure 5.**
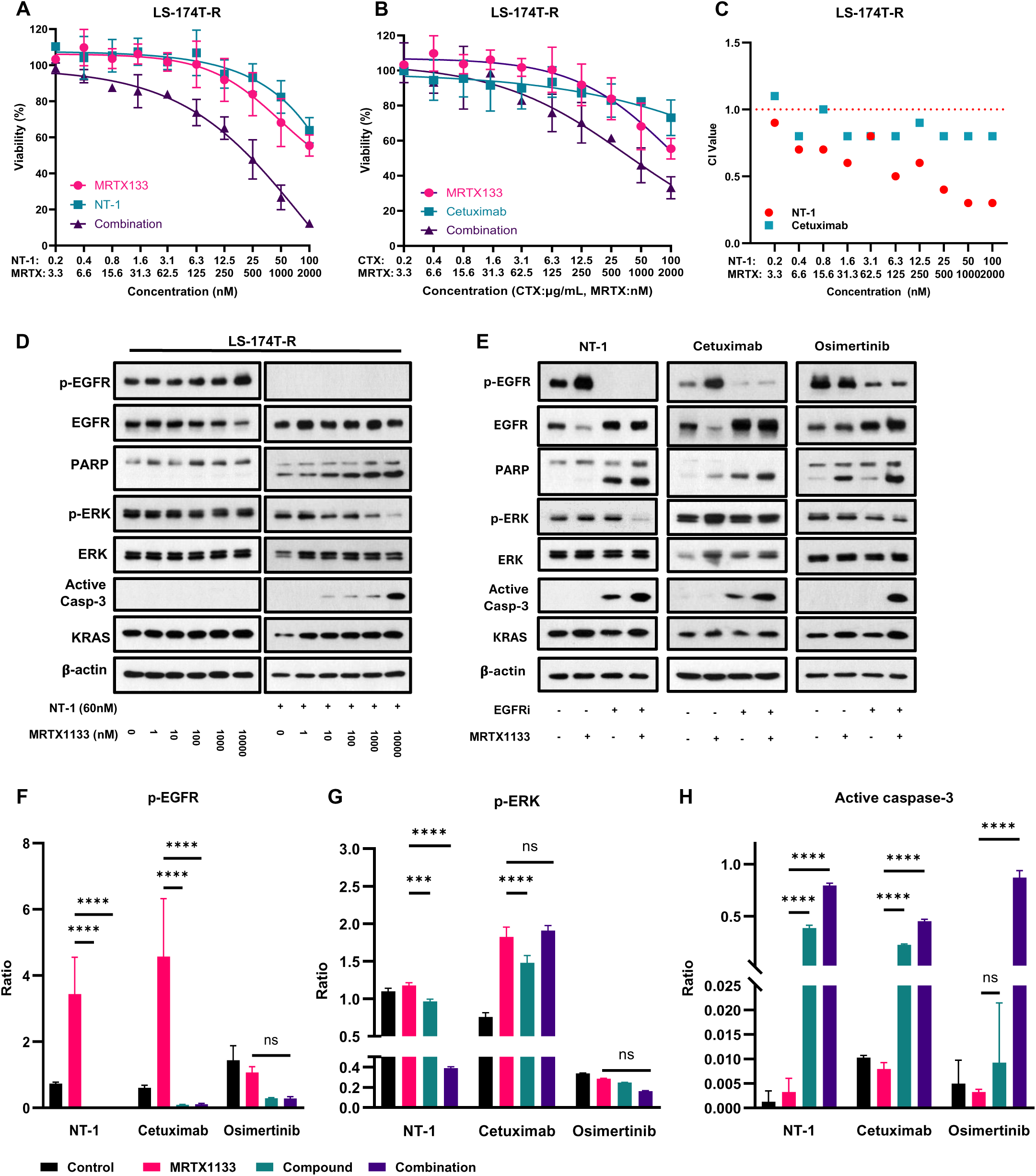
NT-1 synergizes with MRTX1133 to strongly suppress the EGFR/MAPK pathway. **A,B.** Dose-response curves in resistant cell line treated with (A) NT-1 and (B) Cetuximab, alone or in combination. **C.** Combination Index of NT-1 (nM) in comparison with Cetuximab (µg/mL) in LS-174T-R cell line at different concentrations, showing more synergy in NT-1 + MRTX1133. **D**. Immunoblot data of LS-174T-R cell line treated with MRTX alone (left) and in combination with NT-1 at 60nM (right), focusing on downstream MAPK pathway proteins. **E.** Immunoblot analysis of NT-1 (60nM) in combination with MRTX1133 (1μM), compared to Cetuximab (100µg/mL) and Osimertinib (1µM), evaluating downstream MAPK pathway proteins. **F,H.** Quantification of p-EGFR (F), p-ERK (G), and cleaved caspase 3 (H) levels with MRTX1133, NT-1, Cetuximab, and Osimertinib monotherapy, and in combination. Statistical analysis was performed using one-way ANOVA with multiple comparisons (****p<0.0001). Data presented as mean ± SD.

This was further characterized in an immunoblot of Cetuximab (100 µg/mL), Osimertinib (1 µM), and NT-1 (60 nM) as monotherapy and in combination treatment with MRTX1133 (1 µM) (**Fig. 5E**). While monotherapy with MRTX1133 induced phospho-EGFR upregulation, cells co-treated with NT-1 failed to activate EGFR, indicating effective suppression of the feedback loop. Cetuximab and Osimertinib combination treatments showed decreased p-EGFR and increased PARP and Caspase 3 expression, but they were less effective compared to NT-1. The suppression of p-EGFR showed a significant decrease in NT-1 monotherapy and in combination with MRTX1133 when compared to MRTX1133 alone (p < 0.0001), and this similar result was observed with Cetuximab, but not with Osimertinib (**Fig. 5F**). This finding validates the previous data on Osimertinib, which is approved for mutant EGFR, not wild-type EGFR ^25^. Therefore, pharmacological inhibition of EGFR by NT-1 sensitizes the cells to the KRAS **^G12D^** inhibitor and effectively prevents the signaling feedback loop, resulting in more complete suppression of MAPK signaling. Looking at the downstream pathway target of p-ERK, NT-1 monotherapy and combination (p < 0.001) successfully inhibit p-ERK compared to Cetuximab and Osimertinib (**Fig. 5G**). Cetuximab treatment alone seemed to lead to p-ERK suppression, but this suppression was not maintained in the combination treatment, possibly due to differences in the mechanisms of action between antibody vs. small molecule inhibitors. Finally, active caspase-3 activity was quantified to show an increase in activity, with all compounds showing increased apoptosis in combination treatment for 24 hours (**Fig. 5H**). Overall, NT-1 appears to be the most effective inhibitor of wildtype EGFR signaling, and when combined with MRTX1133 can further inhibit downstream MAPK pathways, potentially reducing resistance mechanisms.

### VI. NT-1 synergizes and overcomes MRTX1133 resistance in Patient Derived Organoids (PDO) models

To further validate the potent synergy between NT-1 and MRTX1133 in overcoming resistance to MRTX1133, we utilized both parental and resistant patient-derived organoid (PDO) models. These PDOs closely resemble the tumor epithelium both phenotypically and genetically, providing a more translational model for study ^26^. The mutational landscape of the KRAS G12D organoid revealed a G12D allele fraction closely aligned with that of the LS174T-P cell line, approximately 50%. Additionally, concurrent oncogenic alterations were identified, including mutations in SOS1, ATM, and APC (**Supplementary Fig. S6A**). ^27^ Similar to the LS-174T parental protocol for the KRAS **^G12D^** cell line, an NCI organoid model was treated with eight repeated cycles MRTX1133 until the partial loss of organoid integrity was observed. Organoids that survived the 8 cycles of treatment were called G12D-PDO-R (MRTX1133 resistant) (**Fig. 6A**). The resistant organoids were maintained in media containing 1µM of MRTX1133. The growth rate and size of the resistant organoids exceeded the parental organoids (**Fig. 6B**). Genomic sequencing of cancer driver genes in the resistant organoids revealed that KRAS ^G12D^ mutation was retained, with no additional new mutations detected (**Supplementary Fig. S6A**). With the parental and resistant MRTX1133 resistant organoids ready, we tested the combination of NT-1 and MRTX1133 *in vitro* to determine if the heterogeneity of CRC PDO models could be overcome by the top synergistic compound (**Fig. 6C, Supplementary Fig. S6B**). Based on dose response curves and synergy assessment, we observed that the combination therapy works at low concentrations (∼10nM) with an average CI value of 0.5. There is a strong synergistic effect as well as efficacy in monotherapy, which is notable because CRC is not driven by EGFR mutations. This is further characterized when comparing NT-1 alone and in combination in the parental and resistant PDO models (**Fig. 6D**) The monotherapy and combination therapy showed significantly decreased cell viability over 5 days compared to MRTX1133 alone, especially in the G12D-PDO-R model (p<0.0001). This was further corroborated by IHC (**Fig. 6E**) analysis of p-EGFR suppression and other downstream pathways. A microscopic view of the organoids undergoing treatment (**Supplementary Fig S6C.**) showed noticeable apoptosis and was further corroborated in the immunoblot (**Fig. 6F**) after just 72 hours of treatment, the organoids exhibit a rapid synergistic inhibitory effect at low concentrations. In the G12D-PDO-R model, combination treatment resulted in a significant decrease in p-EGFR (**Fig. 6H**, p<0.0001) and its downstream target p-ERK (**Fig. 6G**, p<0.001), along with an increase in c-PARP (**Fig 6I**, p<0.00001). This led to downstream suppression and increased apoptosis, sensitizing the organoid to treatment that had become resistant to MRTX1133 alone. These findings strongly indicate that NT-1 combined with MRTX1133 treatment can overcome acquired resistance mechanisms in the treatment of KRAS^G12D^ mutant CRC.

**Figure 6.**
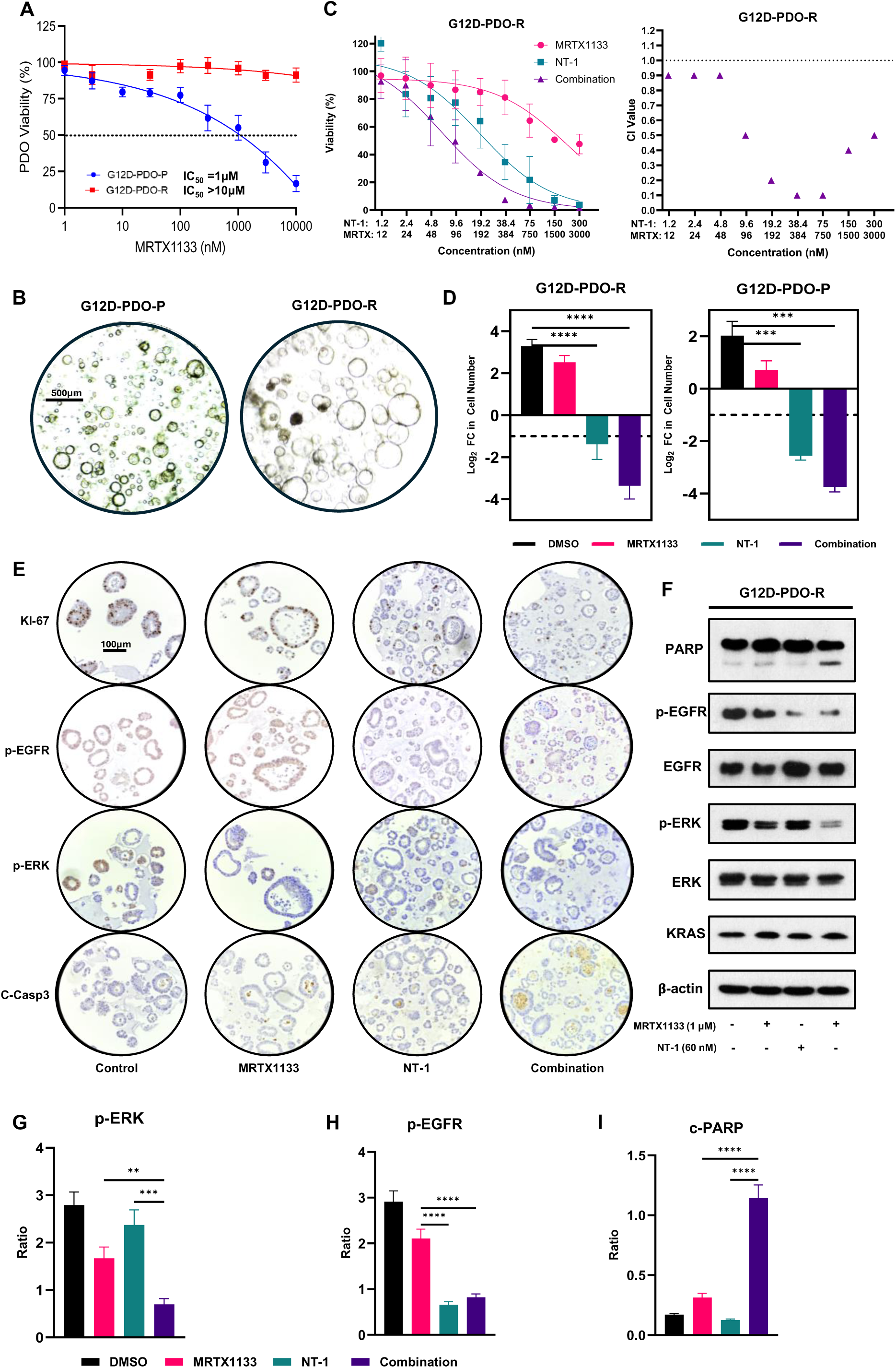
NT-1 and MRTX1133 strongly synergize to suppress tumor growth and EGFR/MAPK signaling in a patient-derived organoid with acquired MRTX1133 resistance. **A.** Proliferation assay for MRTX1133 resistant KRAS^G12D^ tumor organoid after treatment with the indicated compounds for 5 days. **B**. Morphology of parental (left) and resistant (right) organoid model. **C.** Dose-response data with NT-1 (starting at 300nM) in combination with MRTX1133 (starting at 3µM) in the G12D-PDO-resistant organoid, along with respective combination index (CI) values for each concentration. **D.** Proliferation assay over 5 days in G12D-PDO-P treated with DMSO, MRTX1133 (3µM), NT-1 (300nM), or combination. Graph reflects the relative change in cell number (log2 fold scale to best visualize loss of cells) compared with day 0. **E**. Immunohistochemistry (IHC) staining of G12D-PDO-R treated with specified conditions for 72 hours. **F.** Immunoblot data of G12D-PDO-R treated with MRTX (1µM) and NT-1 (60nM) for 72 hours, focusing on various downstream MAPK pathway proteins. **G-I.** Quantification of p-ERK (G), p-EGFR (H), and cleaved PARP (I) levels with MRTX1133 and NT-1 individually, and in combination. Statistical analysis was performed using one-way ANOVA with multiple comparisons (****p<0.0001). Data presented as mean ± SD.

## DISCUSSION

Our study provides significant insights into combination therapies with MRTX1133 to improve CRC outcomes. Recognizing that synergy in molecular combinations is rare and varies across tissues and molecular backgrounds ^28^, our approach underscores the importance of evaluating both combination potency and efficacy. We identified compounds that synergize with the KRAS ^G12D^ inhibitor MRTX1133 in both parental and resistant CRC cell lines through high-throughput screening. By utilizing a screening concentration at 10nM with one of the largest kinase inhibitor library screens in cancer ^28,29,30^, we aimed to identify combinations that enable dose reduction, improved efficacy, or both, relative to single-agent treatments. Importantly, combinations that were not cytotoxic in the monotherapy conditions were prioritized to avoid off-target and non-specific cytotoxicity. By further characterizing the 187 primary hits, we identified promising kinase inhibitors that target proteins such as Akt, Raf, and EGFR. MRTX1133 was present in our library and emerged as an additive, but not synergistic molecule in our screen. This finding serves as an internal control for our screen. When comparing the overall profile of our primary hits, similar targets and proteins were found through a recent CRISPR-Cas9 synthetic lethal screen, which identified EGFR and PLK1 as critical targets for co-treatment with MRTX1133 in LS174T cells, aligning with our screen’s findings that EGFR inhibitors synergize effectively with MRTX1133 ^13^.

Our primary hits were categorized into 16 groups based on the expected outcomes of the four screening groups (**Supplementary** Fig.2C.). Categories 2 and 5 targeted general KRAS synergizers, while categories 3, 6, and 7 targeted resistant-selective KRAS synergizers and compounds against resistant cancer cells. Focusing on differences between parental and resistant cell lines, we plotted the number of hit compounds from each group against various biological pathways (**Supplementary** Fig.2D.). Notably, the number of compounds in the MAPK/ERK and JAK/STAT pathways increased in resistant versus parental cell lines, indicating a potential reliance on these pathways for survival (**Fig. 2D**). To ensure selective cytotoxicity in the presence of MRTX1133, our secondary synergizer hit cutoff required compounds to demonstrate at least a 30% difference in inhibition between monotherapy and the MRTX1133 combination. Using this stringent threshold, 18 compounds were identified that demonstrate synergy with MRTX1133 in both parental and resistant cell lines. The key targets for driving synergy included EGFR, Ser/Thr, p38, and Raf which exhibited above 50% synergy between monotherapy and combination treatments (**Fig. 2F**). Interestingly, the hit molecules fell into two main categories: vertical inhibitors of the RTK-RAS-RAF-MEK pathway and cell cycle inhibitors.

Building on these findings, particularly the strong synergy observed with EGFR and related kinase inhibitors, we explored combinations of MRTX1133 with multiple promising kinase inhibitors to address resistance and enhance therapeutic efficacy. EGFR inhibitors, NT-1, and its analogs exhibited the most striking synergy, evident across both parental and resistant models, highlighting their potential in overcoming MRTX1133 resistance. The combination effectively downregulated the MAPK pathway, a critical signaling cascade in CRC progression. These findings are similar to results in pancreatic ductal adenocarcinoma (PDAC) models, where MRTX1133 demonstrated specificity and effectiveness *in vitro* ^31,32^. The overexpression of phosphorylated EGFR following MRTX1133 treatment suggests that combining MRTX1133 with inhibitors targeting these receptors can enhance anticancer activity. Interestingly, the low nanomolar potency of NT-1 outperformed cetuximab and Gefitinib *in vitro*, showing significantly greater synergy in combination treatments. Cetuximab combined with MRTX1133 in CRC cell lines from other groups demonstrated a ZIP score of ∼17 ^13,33^, whereas NT-1 displayed significantly higher potency, paving the way for its exploration as a less toxic alternative to other wild-type EGFR treatments. The overexpression of phosphorylated EGFR following MRTX1133 treatment suggests a compensatory feedback loop, but additional pathways, such as c-Met and Raf signaling, also appear to contribute to the response. Similarly, studies involving osimertinib, gefitinib, and other EGFR inhibitors have shown promising synergy with KRAS-targeted therapies^17,20,34^. Notably, NT-1 exhibited superior potency in our study compared to cetuximab and gefitinib, reinforcing its potential as a leading candidate for further exploration.

Our study highlights the challenge of acquired resistance to KRAS inhibitors. MRTX1133 is a relatively new inhibitor with limited studies on resistance mechanisms ^10,24,33^. We identified a novel KRAS mutation, KRAS ^E62D^, which could weaken MRTX1133’s potency. While approximately 90% of KRAS mutations occur within codons 12 and 13, there is minimal data on mutations like KRAS ^E62D^. Based on the design of MRTX1133 and the expected interaction between the pyrrolizidine moiety and the E62 carboxylate group, it is possible that a KRAS ^E62D^ mutation leads to reduced binding affinity of MRTX1133 to KRAS protein^12^. As further support for this theory, a prior study by Choi *et al* that used deep mutational scanning analysis indicated that KRAS ^E62D^ is associated with resistance against MRTX1133 ^35^. This mutation may represent a novel resistance mechanism specific to KRAS G12D inhibitors, warranting further investigation. Additionally, resistance-associated pathways, such as Src and PLK1, emerged from our analysis, suggesting alternative mechanisms beyond direct KRAS mutations. Strikingly, our detection of Src aligns with a recent study that showed the SRC-JUN pathway, activated via RAF1-MEK-ERK and potentially JNK, drives G12C inhibitor resistance, raising the possibility that analogous MAPK-dependent mechanisms underlies G12D inhibitor resistance ^36^. Understanding these resistance mechanisms will be crucial for designing effective combination strategies. Our screening process identified numerous targets and pathways relevant to KRAS ^G12D^ inhibition. The hit compounds were categorized based on their effects in parental and resistant models, with significant enrichment observed in the MAPK/ERK and JAK/STAT pathways. The compounds that demonstrated synergy with MRTX1133 were mapped to their respective targets and pathways, reinforcing the importance of dual-targeting strategies. Given the observed reliance on MAPK signaling in resistant models, our study underscores the potential of targeting multiple nodes within this pathway to enhance therapeutic outcomes. We identified and confirmed EGFR inhibition with compounds like Gefitinib and MRTX1133 are synergistic as well, but the potency of NT-1 is significantly greater. This could pave the way for exploring other combinations with NT-1, providing a less toxic alternative to other wild type EGFR treatments. In conclusion, our study highlights the value of exploring synergistic drug combinations to overcome resistance mechanisms in CRC. With appropriate consideration of potential toxicity and molecular profiling of patient tumors, these combinations could offer more effective and personalized therapeutic options for patients with KRAS ^G12D^ CRC.

Limitations to our study must be acknowledged. Our screening process identified numerous targets and pathways, which unfortunately means that this study does not fully represent the breadth and depth of the screen. This leaves us with important information that can be leveraged in future studies. Despite having two distinct models resistant to MRTX1133, it is crucial to further characterize the transcriptional profiles across multiple models. This will help us gain a deeper understanding of the most likely mechanisms driving resistance. Additionally, further studies are needed to evaluate the efficacy of these combinations in murine models. Our data indicate that combined inhibition enhances the initial response and may prolong the durability of the response. Hence, identifying a comprehensive list of resistance-promoting factors is necessary. This information will provide actionable predictive biomarkers, allowing us to personalize combinations with KRAS inhibitors for patients in forthcoming clinical trials. We also observed early signs of genetic resistance mechanisms to MRTX1133, necessitating further investigation to understand their prevalence and identify strategies to effectively restore treatment response. It will be intriguing to study how clinical resistance to upcoming pan-KRAS inhibitors develops, as this will inform future therapeutic approaches.

In conclusion, our study provides a holistic and unbiased screening of a diverse kinase library leading to the identification and discovery of Osimertinib analogs synergizing with MRTX113 in KRAS **^G12D^**mutant colorectal cancer to overcome resistance to MRTX1133 monotherapy. We used patient-derived organoid models to confirm the synergy between the EGFR inhibitor NT-1 and MRTX1133, providing strong rationale for clinical testing of this combination in patients with KRAS **^G12D^**mutant colorectal cancer.

## MATERIALS AND METHODS

### Cell Viability Assay

Cell viability was assessed using the CellTiterGlo 2.0 Luminescent Cell Viability Assay (Promega). Cells were seeded in 96-well plates at a density of 3,000 cells per well or 384-well plates at a density of 1,000 cells per well and incubated overnight at 37°C in a 5% CO₂ atmosphere to allow for adherence. Post-treatment with experimental compounds for 72 hours, 20-100 µL of CellTiter-Glo reagent was added to each well, followed by gentle mixing on an orbital shaker for 2 minutes. After a 10-minute incubation at room temperature, luminescence was measured using a relative luminescence units (RLUs) from treated wells were normalized to control wells to determine cell viability. IC_50_ values were calculated by GraphPad Prism (RRID: SCR_002798) using 3-parameter dose-response model.

### Cell Lines and Cell Culture

LS513 (RRID: CVCL_1386), LS-174T (RRID: CVCL_1384), HCT116 (RRID: CVCL_0291), RKO (RRID: CVCL_0504), and HT29 (RRID: CVCL_0320) human colon cancer cell lines were purchased from American Type Culture Collection (ATCC, Manassas, VA, USA). LS513 cells were cultured in RPMI1640 (FUJIFILM Wako Pure Chemical Corporation, Osaka, Japan) supplemented with 10% fetal bovine serum and 1% P/S. LS-174T cells were cultured in Dulbecco’s modified Eagles medium (DMEM; FUJIFILM Wako Pure Chemical Corporation) supplemented with 10% fetal bovine serum and 1% P/S. All cell lines were incubated at 37°C and 5% CO_2_. Cell lines were authenticated every 6 months while in use. Cell lines were monitored every 6 months for Mycoplasma infection and treated when necessary.

### Establishment of MRTX1133 Resistant Cells

MRTX1133-resistant cells were established by exposure of LS-174T cells to increasing concentrations of MRTX1133. The initial concentration of MRTX1133 was 10 nM. When the cells adapted to the drug, the concentration of MRTX1133 was gradually increased by 1.5–2 times every week to a final concentration of 1 μM.

### Cell Seeding and Screening

Freshly trypsinized colorectal cell lines were seeded into black CELLSTAR® uCLEAR® 384 well plates (Greiner, Monroe, NC, USA) with 1000 cells per well. After cell seeding, the cells were incubated overnight to adhere to the bottom of the plate before drugs were added. The kinase inhibitor library was purchased from a vendor (MedChemExpress, RRID: SCR_025062). All kinase inhibitors were tested at 10 nM, and, for selected hit compounds, a concentration range of 1 - 1,000 nM was tested. Cells were incubated for 72 h with small molecules in media at 37 °C, 5% CO_2_ before viability assay read out.

### Reverse-phase protein array (RPPA)

LS-174T-P and LS-174T-R cells were treated with MRTX1144 (1000nM) for 24 h, the cells were harvested, and protein lysates were prepared. RPPA was performed by the Functional Proteomics RPPA Core Facility at the MD Anderson Cancer Center (RRID: SCR_016649) for analysis.

### Synergy determination with the SynergyFinder method

CRC Cell lines were seeded at a density of 0.05 × 10^6^ cells/well in 384-well black clear plates (Greiner) and were further treated as described above. Synergy scoring was determined using the “inhibition readout” (calculated as “100 - Cell Viability” on the online SynergyFinder software, RRID: SCR_019318) and implementing the ZIP calculation method. The matrix started with 1 µM in the hit compounds and 300 nM or 3,000 nM in the MRTX1133 starting concentration with a 3-fold decrease in the parental and resistant, respectively. The cell viability results were input into synergyfinge3.0 to determine synergistic potential. The ZIP synergy score ^14^ was used to evaluate the synergisms between compound and MRTX1133. Scores above 10 are synergistic, between - 10 and 10 are additive, and below -10 is considered antagonistic.

### Western Immunoblot

Cell lysates were harvested using a lysis buffer supplemented with protease and phosphatase inhibitors. Proteins from each sample (10–50 µg) were separated on 4—20% mini-PROTEAN TGX gels and transferred to nitrocellulose membranes (Bio-Rad, Hercules, CA, USA). After blocking in 5% milk with Tris-buffered saline with Tween (TBST) buffer, the membranes were probed with primary antibodies overnight at 4 °C. The membranes were washed three times in TBST and incubated with secondary antibodies for 1 h at room temperature. Image acquisition and band intensity quantification were performed using an Odyssey infrared imaging system (LI-COR Biosciences, LincoIn, USA) and Image J software (NIH, Bethesda, USA, RRID: SCR_003070), respectively. The following primary antibodies were obtained from Cell Signaling Technology and used at a dilution of 1:1000: anti-EGFR (#4267, RRID: AB_2895042), anti-phospho-EGFRY1068 (#3777, RRID: AB_2096270), anti-ERK1/2 (#9102, RRID: AB_330744), anti-phosphoERK1/2 (#4370, RRID: AB_2315112), anti-AKT (#9272, RRID: AB_329827), anti-phospho-AKTS473 (#4060, RRID: AB_2315049), and anti-PARP (#9542, RRID: AB_2160739).

### Patient-derived organoids

In brief, frozen aliquots of PDO tissue were obtained from NCI Patient-Derived Models Repository were obtained, thawed, and re-established as PDOs in our lab. Thawed tissues were embedded in Matrigel and cultured in complete PDO media which was supplemented with WRN conditioned media as well as ROCK inhibitor. All PDOs used were passaged at least twice and growing with a doubling time of 3-10 days, prior to use in experiments. Brightfield micro-photographs were obtained using the Echo Revolve 4 (RRID: SCR_026523).

### Genomic sequencing

Cancer-focused mutation analysis was performed using the Ion Torrent AmpliSeq Colon Cancer Panel (AmpliSeq Library 96LV Kit 2.0, ThermoFisher). Library preparation, quality control, sequencing, and data analysis were performed as per manufacturer instructions.

### Immunohistochemistry

PDOs were grown and treated, then isolated from Matrigel domes using Cell Recovery Solution (Corning #354253). PDOs were fixed with 4% PFA, embedded in 1% agarose, then processed into FFPE blocks. 5 µm thick sections were subjected to standard IHC. Visualization was with ImmPACT DAB Substrate Kit (Vector Laboratories #SK-4105). Brightfield micro-photographs were obtained using the Echo Revolve 4 (RRID: SCR_026523).

### Kinase Assay

The “HotSpot” assay platform was performed by Reaction Biology using 10 different concentrations of NT-1 across a select panel of recombinant kinases ^37^. ATP concentration of 10 µM was used for the assay. Recombinant proteins included: EGFR (Invitrogen, Cat# PR7295B), EGFR T790M (Invitrogen, Cat# PR8052A), IGF1R (Invitrogen, Cat# PV3250), c-SRC (Invitrogen, Cat# PR4336E), and PLK1 (BPS, Cat# 40033). Briefly, specific kinase / substrate pairs along with required cofactors were prepared in reaction buffer: 20 mM Hepes pH 7.5, 10 mM MgCl_2_, 1 mM EGTA, 0.02% Brij35, 0.02 mg/mL BSA, 0.1 mM Na_3_VO_4_, 2 mM DTT, 1% DMSO, Compounds were delivered into the reaction, followed ∼ 20 minutes later by addition of a mixture of ATP (Sigma, St. Louis MO) and 33P ATP (Perkin Elmer, Waltham MA) to a final concentration of 10 μM. Reactions were carried out at room temperature for 120 min, followed by spotting of the reactions onto P81 ion exchange filter paper (Whatman Inc., Piscataway, NJ). Unbound phosphate was removed by extensive washing of filters in 0.75% phosphoric acid. After subtraction of background derived from control reactions containing inactive enzyme, kinase activity data was expressed as the percent remaining kinase activity in test samples compared to vehicle (dimethyl sulfoxide) reactions. IC_50_ values and curve fits were obtained using GraphPad Prism Software (RRID: SCR_002798).

## List of Supplementary Materials

Fig. S1 to S6

Table S1

Table S2

## Acknowledgments

1. S. Kitamura and C. Kuang are grateful to Einstein-Montefiore for the support to start the lab. Drs. Juan Du and Doctor Y. Goldstein (Montefiore Einstein, Dept. of Pathology) assisted with performing IonTorrent sequencing. FFPE specimen processing, slide cutting, and H&E staining was conducted by Laura Ramkissoon of the Albert Einstein College of Medicine Histology and Comparative Pathology Facility (P30CA013330). The Functional Proteomics Reverse Phase Protein Array Core was supported in part by The University of Texas MD Anderson Cancer Center (P30CA016672 and R50CA221675).

## Author contributions

Conceptualization: NT, SK, CK

Methodology: SK, CK

Investigation: NT, NW, EN, SK, CK

Visualization: NT, SK, CK

Funding acquisition: EC, SK, CK

Project administration: SK, CK

Supervision: SK, CK

Writing – original draft: NT, SK, CK

Writing – review & editing: NT, NW, SK, CK

## Data and materials availability

All original data will be provided upon reasonable request. Biological materials must have a materials transfer agreement established prior to transfer.

