## Supplementary figures and tables for "High Throughput Screening Identifies Small Molecules that Synergize with MRTX1133 Against Acquired Resistant KRAS ^G12D^ Mutated CRC"

**Supplementary Figure S1.**

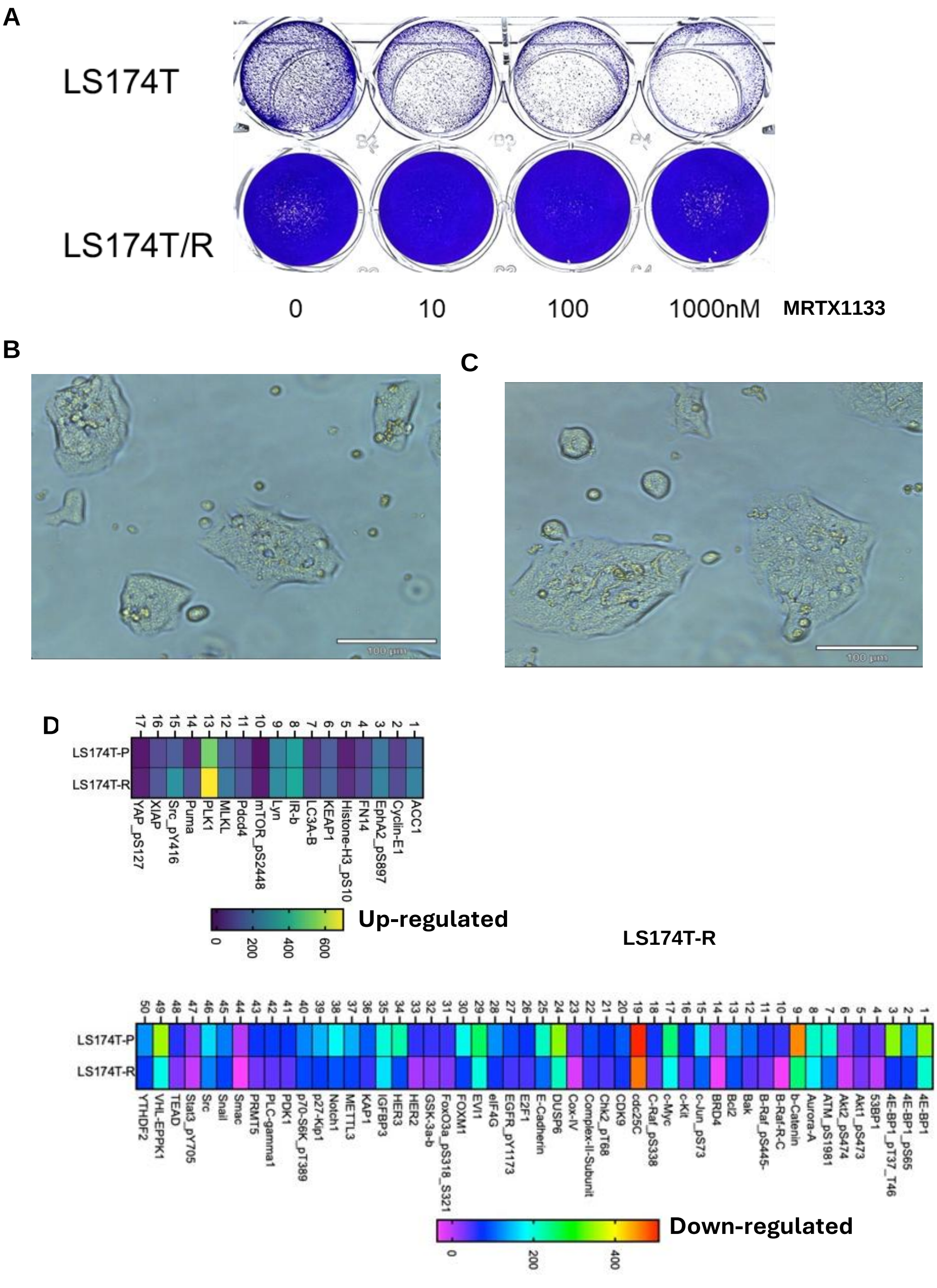


**Supplementary Figure S1.** KRAS G12D resistant cell generation and characterization. **A.** Representative images of LS174T-P and LS-174T-R cells treated with concentration gradients of MRTX1133 for 72hrs. **B,C.** Morphology of LS-174T-P (B) and LS-174T-R (C). **D**. Up regulated and down-regulated genes from RPPA data highlighting cdc25C as most down-regulated.

**Supplementary Figure S2.**

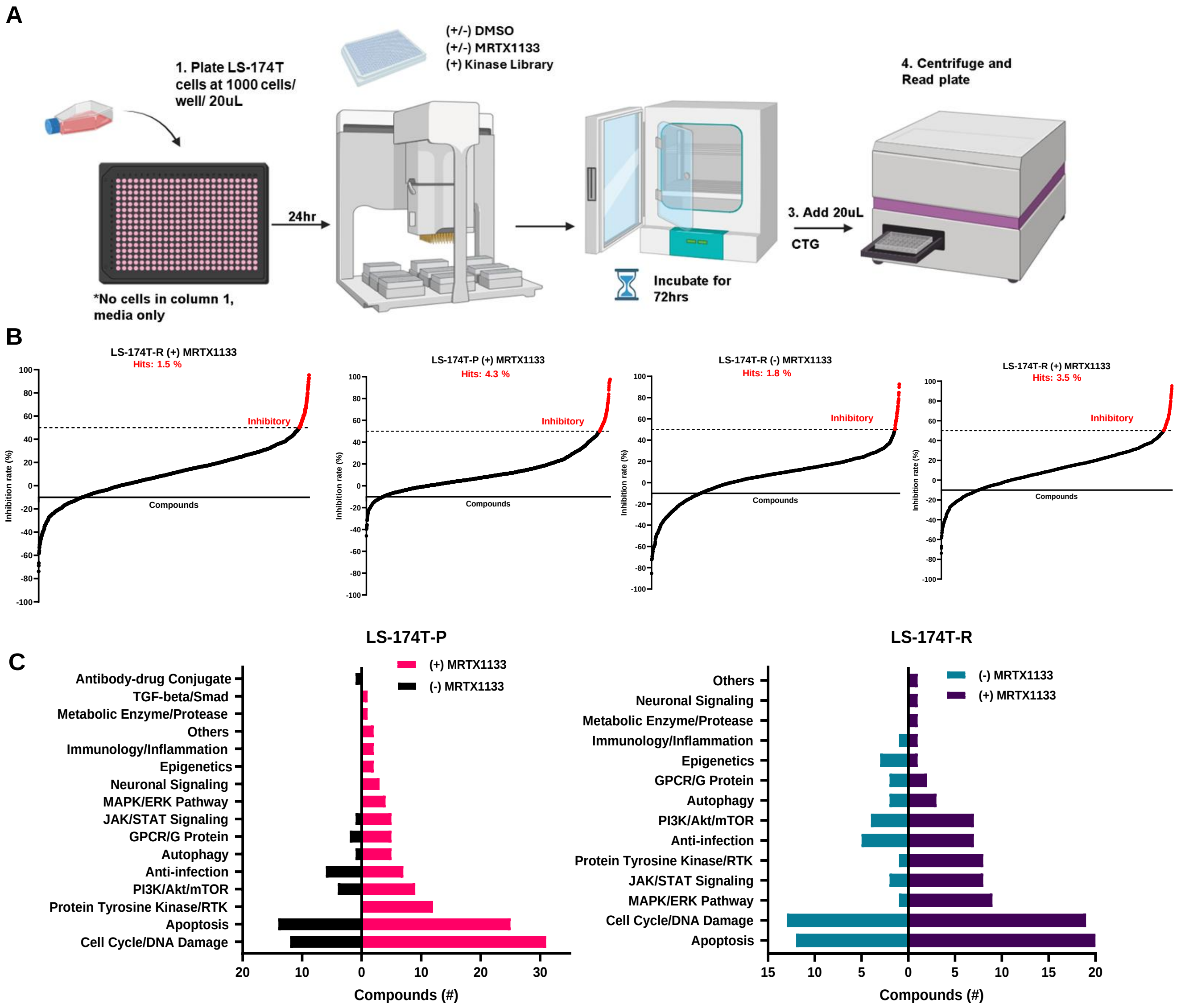


**Supplementary Figure S2.** Screening overview and characterization. **A**. Screening overview schematic. **B**. Inhibition curves for each individual screening condition and the hit percentages. **C**. Complete compounds count of the 187 compounds and their associated group and pathway.

**Supplementary Figure S3.**

A

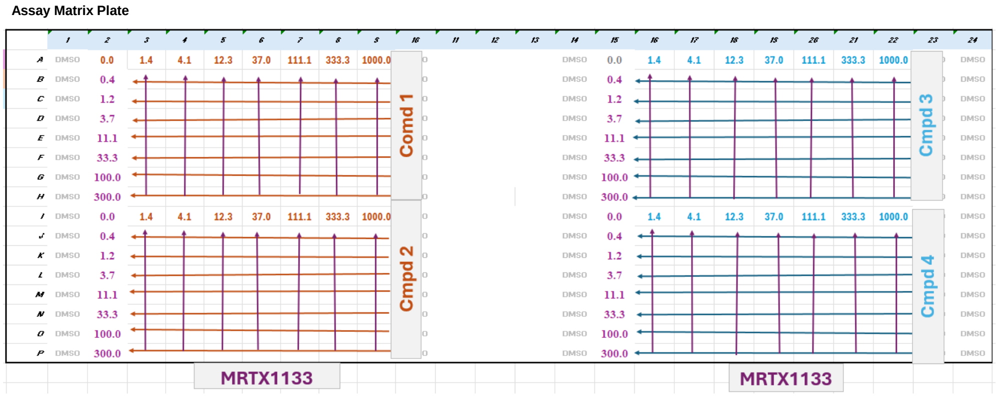


B

| Drug Combination | Cell Line | Synergy Score | Method | CDK Start Conc. |
|---|---|---|---|---|
| CDK8-IN-4 - MRTX1133 | LS-174T-R | 10.432 | ZIP | 100 |
| CDK12-IN-E9 - MRTX1133 | LS-174T-R | 3.559 | ZIP | 10 |
| CDK8-IN-4 - MRTX1133 | LS-174T-P | 1.103 | ZIP | 100 |
| CDK8-IN-4 - MRTX1133 | LS-174T-R | -0.194 | ZIP | 10 |
| CDK12-IN-E9 - MRTX1133 | LS-174T-R | -2.15 | ZIP | 100 |
| CDK12-IN-E9 - MRTX1133 | LS-174T-P | -2.963 | ZIP | 10 |
| CDK8-IN-4 - MRTX1133 | LS-174T-P | -3.771 | ZIP | 10 |
| CDK12-IN-E9 - MRTX1133 | LS-174T-P | -4.839 | ZIP | 100 |

**Supplementary Figure S3.** Matrix hit validation. **A.** Dose-response matrix plate example, can be used multiple times. **B**. Table of decreased CDK inhibitor matrices.

**Supplementary Figure S4.**

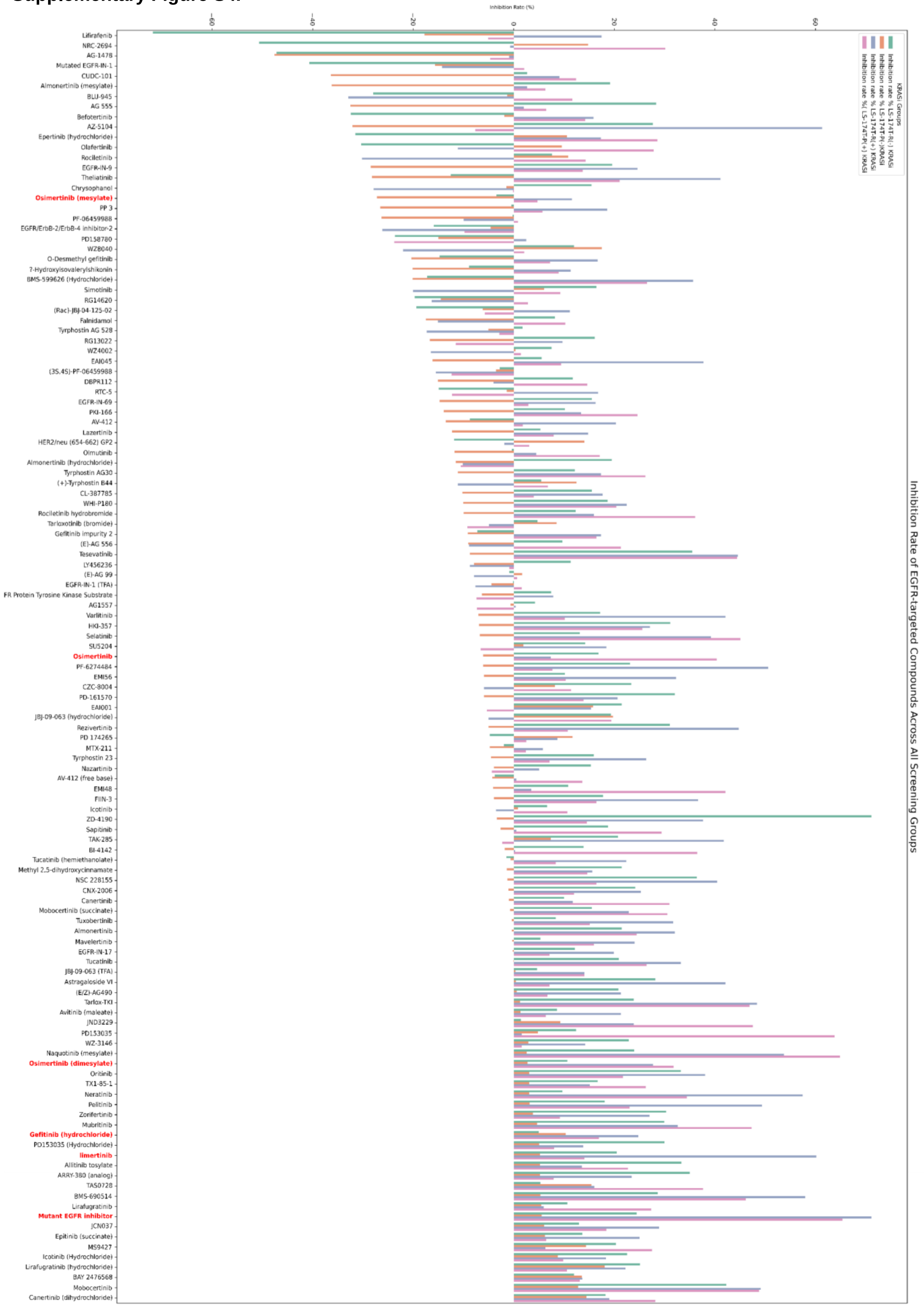

**Supplementary Figure S4.** Inhibition rates of all putative EGFR inhibitors from primary screen.

**Supplementary Figure S5.**

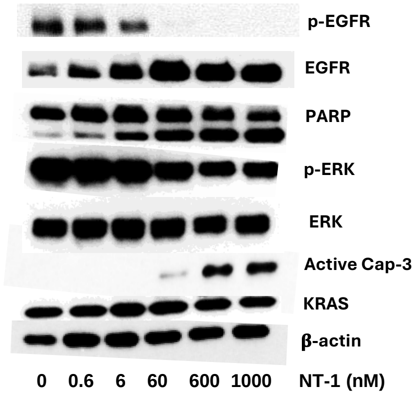


**Supplementary Figure S5.** Immunoblot of monotherapy NT-1 treated in LS-174T-R cell line for 24 hours at escalating concentration. We see p-EGFR suppression at 60nM.

**Supplementary Figure S6.**

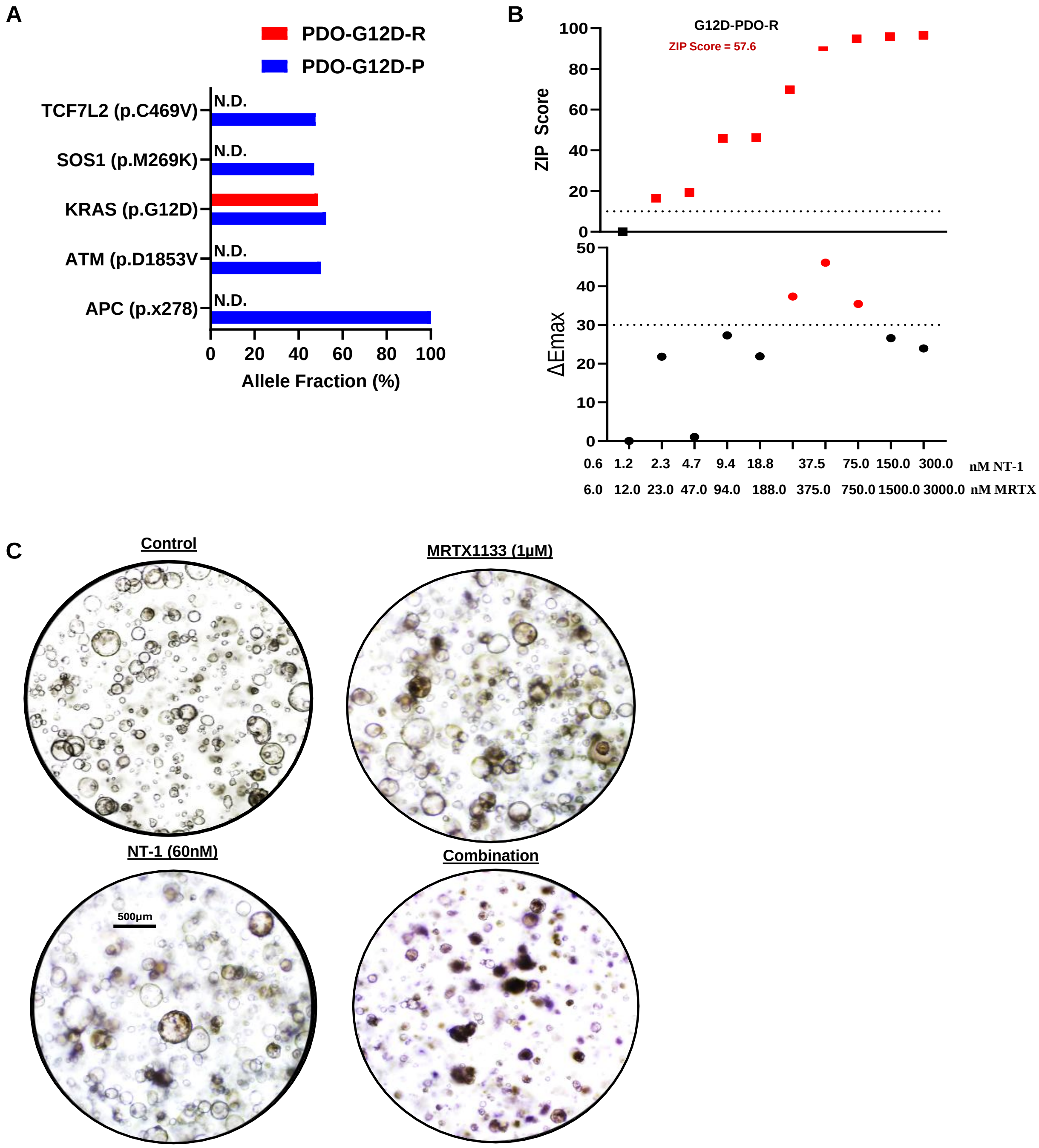


**Supplementary Figure S6.** NT-1 in combination with MRTX1133 in resistant KRAS $^{G12D}$ PDO model. **A.** Mutational profile of G12D-PDO. Parental (P) mutations from NCI PDMR database (ID: 362531). KRAS $^{G12D}$ mutation and lack of EGFR mutation confirmed in Resistant (R) PDO model. N.D.: not determined (not part of IonTorrent sequencing panel). **B.** ZIP and Emax score for NT-1 and MRTX1133 treated in combination of 5 days. **C.** Morphology of G12D-PDO-R treated with MRTX1133, NT-1 individually and in combination for 72 hours, immediately before lysate was collected for immunoblot.

| ProductName | LS-174T-P(-)KRASi | ( LS-174T-P(+) KRASi | LS-174T-R(-) KRASi | LS-174T-R(+) KRASi | Smiles |
|---|---|---|---|---|---|
| (±)-Enitociclib | 14.9 | 51.9 | 30.7 | 46.7 | N=S(CC1=CC(NC2=NC=C(F)C(C3=CC=C(F)C=C3OC)=C2)=NC=C1)(C)=O |
| 1,3-Dicaffeoylquinic acid | 3.8 | 8.5 | 77.0 | 37.2 | O=C([C@@]1(OC(/C=C/C2=CC=C(O)C(O)=C2)=O)C[C@@H](OC(/C=C/C3=CC=C(O)C(O)=C3)=O)[C@H](O)[C@H](O)C1)O |
| 1-Deoxynojirimycin (hydrochloride) | 0.7 | 1.2 | 62.1 | 19.7 | O[C@@H]1[C@@H](CO)NC[C@H](O)[C@H]1O.[H]Cl |
| 8-CPT-Cyclic AMP (sodium) | -3.4 | 62.1 | 16.7 | 29.8 | NC1=NC=NC2=C1N=C(N2[C@@H]3O[C@@]4([H])[C@](OP(OC4)(O[Na])=O)([H])[C@H]3O)SC5=CC=C(Cl)C=C5 |
| A-443654 | 22.1 | 18.5 | 28.8 | 56.4 | N[C@H](COC1=CN=CC(C2=CC3=C(NN=C3C)C=C2)=C1)CC4=CNC5=CC=CC=C54 |
| A-770041 | -7.4 | 66.8 | 20.8 | 37.1 | NC1=C2C(N([C@@H]3CC[C@@H](N4CCN(C(C)=O)CC4)CC3)N=C2C5=CC=C(NC(C6=CC7=C(C=CC=C7)N6C)=O)C(OC)=C5)=NC=N1 |
| Abemaciclib (methanesulfonate) | 24.8 | 52.5 | 29.8 | 40.4 | CC(N1C2=CC(C3=NC(NC4=NC=C(CN5CCN(CC)CC5)C=C4)=NC=C3F)=CC(F)=C2N=C1C)C.CS(=O)(O)=O |
| AEE788 | -3.9 | 54.8 | 5.2 | 33.7 | C[C@H](C1=CC=CC=C1)NC2=C(C=C3C(C=C4)=CC=C4CN5CCN(CC)CC5)C(N3)=NC=N2 |
| Afatinib | -13.7 | 49.3 | 45.0 | 51.1 | O=C(NC1=C(C=C2C(C(NC3=CC(Cl)=C(C=C3)F)=NC=N2)=C1)O[C@H]4CCOC4)/C=C/CN(C)C |
| Afatinib (dimaleate) | 17.1 | 49.4 | 45.1 | 65.2 | O=C(NC1=C(C=C2C(C(NC3=CC(Cl)=C(C=C3)F)=NC=N2)=C1)O[C@H]4CCOC4)/C=C/CN(C)C.O=C(O)/C=C\C(O)=O.O=C(O)/C=C\C(O)=O |
| Agerafenib | -8.1 | 52.4 | 21.5 | 71.3 | O=C(NC1=NOC(C(C)(C)C(F)(F)F)=C1)NC2=CC=C(OC3=C(C=C(OC)C(OC)=C4)C4=NC=N3)=C2 |
| AKI603 | 51.0 | 69.6 | 21.7 | 17.0 | O=[N+]([O-])C1=CC=C(C=C1)NC2=NC(N3CCN(CC3)C)=CC(NC4=NNC(C)=C4)=N2 |
| ALK inhibitor 1 | 3.8 | 28.0 | 4.4 | 56.4 | O=S(C1=C(NC2=C(Br)C=NC(NC3=C(OC)C=C(N4CCN(C)CC4)C=C3)=N2)C=CC=C1)(NC)=O |
| ALK kinase inhibitor-1 | -15.4 | -8.8 | 7.5 | 58.2 | CN(CC1)CCN1C2=CC=C(NC3=NC=C4C(C(C5=C(OC)C=CC(F)=C5)=C(CO)S4)=N3)C(OC(C)C)=C2 |
| ALK2-IN-4 (succinate) | 5.6 | 59.2 | 8.4 | 10.2 | FC(C=C1C(C(C2=C3N(C=C(C=N3)N4CCC(CC4)N5CCN(CC5)C)N=C2)=CC=N1)=C6)=C6OC.O=C(O)CCC(O)=O |
| Alpelisib | 13.3 | 62.4 | 23.4 | 46.6 | CC(N=C(S1)NC(N2CCC[C@H]2C(N)=O)=O)=C1C3=CC(C(C)(C(F)(F)F)C)=NC=C3 |
| ALW-II-41-27 | -1.0 | 4.0 | 78.9 | 46.7 | CC1=C(NC(C2=CC(C3=CC=CS3)=CN=C2)=O)C=C(C(NC4=CC=C(CN5CCN(CC)CC5)C(C(F)(F)F)=C4)=O)C=C1 |
| Amarogentin | 3.8 | 6.6 | 60.5 | 39.8 | C=C[C@H]([C@@]1([H])C(C(OCC1)=O)=CO2)[C@@H]2O[C@@](O[C@H](CO)[C@@H](O)[C@@H]3O)([H])[C@@H]3OC(C4=C(C5=CC(O)=CC=C5)C=C(O)C=C4O)=O |
| AMG 900 | 46.6 | 53.1 | 4.1 | 55.7 | CC1=CSC(C2=NN=C(C3=C2C=CC=C3)NC4=CC=C(C=C4)OC5=C(C=CC=N5)C6=CC=NC(N)=N6)=C1 |
| APS6-45 | 16.8 | 62.3 | 14.9 | 33.3 | O=C(C1=NC=CC(OC2=CC=C(C=C2)NC(NC3=CC(C(F)(C(F)(F)F)C(F)(F)F)=CC=C3F)=O)=C1)NC |
| ARQ 531 | -3.4 | 57.6 | 22.1 | 65.1 | O=C(C1=C(C=C(OC2=CC=CC=C2)C=C1)Cl)C3=CNC4=NC=NC(N[C@@H]5CC[C@@H](CO)OC5)=C34 |
| AS-252424 | 2.3 | 6.1 | 66.3 | 39.9 | O=C(NC/1=O)SC1=C/C2=CC=C(C3=CC=C(F)C=C3O)O2 |
| ASN007 | 27.6 | 10.0 | 26.1 | 55.4 | O=C(C1=CN(C2=NC(NC3CCOCC3)=NC=C2C)C=N1)N[C@@H](C4=CC(F)=CC(Cl)=C4)CN |
| ASP5878 | 2.8 | 41.5 | 18.7 | 51.9 | OCCN1N=CC(NC2=NC=C(OCC3=C(F)C(OC)=CC(OC)=C3F)C=N2)=C1 |
| Atuveciclib S-Enantiomer | 8.0 | 58.4 | 22.7 | 45.3 | N=[S@@](CC1=CC(NC2=NC(C3=CC=C(F)C=C3OC)=NC=N2)=CC=C1)(C)=O |
| Aurora kinase inhibitor-8 | 22.8 | 59.2 | 27.5 | 22.9 | O=C(NC1=CC=C(NC2=NC=C(N3C)C(N(C)C4=CC=C(C)C=C4C3=O)=N2)C=C1)C5=CC=C(NC(CC)=O)C=C5 |
| AZ-23 | -1.6 | 54.6 | 13.5 | 37.6 | CC(OC1=CC(NC2=NC(N[C@H](C3=NC=C(F)C=C3)C)=NC=C2Cl)=NN1)C |
| AZ32 | -3.2 | 2.6 | 55.7 | 30.1 | O=C(NC)C(C=C1)=CC=C1C2=CN=C3C=NC(C4=CC=CC=C4)=CN32 |
| AZ-5104 | -32.1 | -7.6 | 27.6 | 61.3 | C=CC(NC1=CC(NC2=NC=CC(C3=CNC4=C3C=CC=C4)=N2)=C(OC)C=C1N(CCN(C)C)C)=O |
| AZ7550 Mesylate | -2.9 | 53.5 | 19.1 | 47.7 | CN1C=C(C2=NC(NC3=CC(NC(C=C)=O)=C(N(CCNC)C)C=C3OC)=NC=C2)C4=CC=CC=C41.O=S(O)(C)=O.O=S(O)(C)=O.O=S(O)(C)=O |
| AZD3229 | 5.7 | 55.2 | 7.0 | 37.0 | COCCOC1=CC(F)=C2C(N=CN=C2NC3=CC=C(NC(CN4C=C(C(C)C)N=N4)=O)C=C3)=C1 |
| AZD4573 | 93.2 | 95.6 | 84.5 | 87.0 | ClC1=CN=C(NC([C@H]2CCC[C@@H](NC(C)=O)C2)=O)C=C1C3=C(CC(C)(C)C4)N4N=C3 |
| AZD8330 | 41.5 | 51.7 | 48.0 | 64.3 | IC1=CC(F)=C(C=C1)NC(N2C)=C(C=C(C2=O)C)C(NOCCO)=O |
| BAY-3827 | -10.1 | 8.6 | 59.9 | 38.0 | CCC1=CC=CC=C1C(NC2=NNC3=C2C=C(C4C(C#N)=C(C)N(C)C(C)=C4C#N)C(F)=C3C)=O |
| BEBT-908 | 85.8 | 94.1 | 53.8 | 67.1 | CN(CC1=CC(N=C(C2=CN=C(NC)C=C2)N=C3N4CCOCC4)=C3S1)C5=NC=C(C(NO)=O)C=N5 |
| BGT226 (maleate) | 64.9 | 78.0 | 26.4 | 84.6 | COC1=CC=C(C=N1)C2=CC=C(C3=C2)N=CC(N4C)=C3N(C4=O)C5=CC=C(C(C(F)(F)F)=C5)N6CCNCC6.OC(/C=C\C(O)=O)=O |
| BI 2536 | 79.5 | 87.9 | 29.8 | 75.9 | O=C1[C@H](N(C2=C(N1C)C=NC(NC3=CC=C(C(NC4CCN(C)CC4)=O)C=C3OC)=N2)C5CCCC5)CC |
| BI-D1870 | -6.7 | 1.0 | 58.0 | 14.7 | OC1=C(F)C=C(NC2=NC(N(CCC(C)C)C(C)C(N3C)=O)=C3C=N2)C=C1F |
| BMS-690514 | 5.3 | 46.1 | 28.6 | 58.0 | O[C@@H]1CN(CC[C@H]1N)CC2=C3C(NC4=CC=CC(OC)=C4)=NC=NN3C=C2 |
| B-Raf IN 1 | 17.6 | -7.8 | 75.9 | 26.4 | O=C(NC1=CC=CC(C2=CC=NC3=C(C4=CC=C(CN(C)C)C=C4)C=NN23)=C1)C5=CC=CC(C(F)(F)F)=C5 |
| Cardamonin | 0.8 | 66.1 | 10.7 | 54.2 | O=C(C1=C(OC)C=C(O)C=C1O)/C=C/C2=CC=CC=C2 |
| Casein Kinase inhibitor A51 | 75.8 | 80.3 | 76.3 | 75.4 | N[C@H]1CC[C@H](NC2=NC=C(Cl)C(C3=C(CC4CC4)N(C)N=C3)=N2)CC1 |
| Casein Kinase inhibitor A86 | 85.5 | 96.0 | 89.8 | 69.2 | N[C@H]1CC[C@H](NC2=NC=C(F)C(C3=C(CC4CC4)N(C)N=C3)=N2)CC1 |
| CCG258747 | -8.3 | 57.1 | 8.1 | 28.4 | O=C(NCC1=NNC2=CC=CC=C12)C(C=C3[C@@](CCNC4)([H])[C@@H]4COC5=CC=C(OCO6)C6=C5)=C(C=C3)F |
| CDK12-IN-3 | 17.7 | 43.3 | 22.5 | 61.0 | CC(C)N1C2=NC(N3CCCC[C@H]3CCO)=NC(NCC4=NC5=C(C(F)=C(F)C=C5)N4)=C2N=C1 |
| CDK12-IN-E9 | 34.5 | 79.8 | 37.7 | 86.8 | C=CC(NC1=CC=CC(NC2=CC(N3[C@H](CCO)CCCC3)=NC4=C(CC)C=NN24)=C1)=O |
| CDK7-IN-2 | 34.0 | 68.9 | 23.1 | 30.2 | O=C(N1CCC(NC2=CC(C)=NC3=C(C(C)C)C=NN23)CC1)O[C@@H]4CN(C(/C=C/CN(C)C)=O)CC4 |
| CDK8-IN-4 | 44.5 | 81.7 | 38.2 | 72.3 | OC[C@@H](C1=CC=CC=C1)NC2=CC(C3=CC4=C(NN=C4)C=C3)=CN=C2 |
| CDK9-IN-13 | 43.3 | 95.0 | 55.6 | 47.3 | CC(C)N1C=C(C2=CC=NC3=C2C=C(C4CCN(CC(N5CCCCC5)=O)CC4)N3)C=CC1=O |
| CDK9-IN-15 | 0.0 | 56.9 | 8.6 | 17.3 | OC1=C2C=CC=C(NC3=C(C=CS4)C4=NC=N3)C2=CC=C1 |
| Cenisertib | 50.7 | 80.1 | 58.5 | 79.1 | [H][C@@]1([C@H]([C@H]2C(N)=O)NC3=NC(NC4=CC=C(C(C)=C4)N5CCN(CC5)C)=NC=C3F)C[C@]2(C=C1)[H] |
| CEP-40783 | -33.8 | 7.4 | -10.1 | 58.7 | O=C(C1=CN(C(C)C)C(N(C2=CC=C(F)C=C2)C1=O)=O)NC3=CC=C(OC4=CC=NC5=CC(OC)=C(OC)C=C45)C(F)=C3 |
| Cevidoplenib | -21.1 | 50.0 | 18.7 | 34.4 | O=C(C1CC1)C2=CN(C)C3=C2C=C(NC4=NC=CC(N5N=C(C)C(CN6OC[C@@H](O)C6)=C5)=N4)C=C3 |
| Copanlisib (dihydrochloride) | 48.5 | 56.7 | 51.2 | 87.2 | O=C(C1=CN=C(N)N=C1)NC2=NC3=C(OC)C(OCCCN4CCOCC4)=CC=C3C5=NCCN25.[H]Cl.[H]Cl |
| CP-547632 | -13.8 | 56.8 | 12.6 | 39.8 | O=C(NC1=C(C(N)=O)C(OCC2=C(F)C=C(Br)C=C2F)=NS1)NCCCCN3CCCC3 |
| Crizotinib | -1.8 | 56.9 | 15.6 | 43.5 | ClC1=C(F)C=CC(Cl)=C1[C@H](OC2=CC(C3=CN(N=C3)C4CCNCC4)=CN=C2N)C |
| CTX-0294885 | 56.5 | 74.4 | 37.0 | 77.0 | O=C(NC)C1=CC=CC=C1NC2=NC(NC3=CC=C(N4CCNCC4)C=C3)=NC=C2Cl |
| CTX-0294885 (hydrochloride) | 74.1 | 75.9 | 67.2 | 52.1 | [H]Cl.O=C(NC)C1=CC=CC=C1NC2=NC(NC3=CC=C(N4CCNCC4)C=C3)=NC=C2Cl |
| Cu(II)GTSM | 49.4 | 78.6 | 41.9 | 83.5 | CNC1=N[N]2=CC=[N]3N=C(NC)[S-][Cu+2]23[S-]1 |
| Cucurbitacin E | 80.5 | 75.9 | 84.7 | 63.2 | O[C@H](C1)[C@@]([C@@](C)(O)C(/C=C/C(OC(C)=O)(C)C)=O)([H])[C@](C2)(C)[C@]1(C)[C@]3([H])CC=C4C(C)(C)C(C(O)=C[C@@]4([H])[C@]3(C)C2=O)=O |
| Dactolisib (Tosylate) | 37.3 | 55.7 | 15.2 | -10.2 | CN(C1=C2C3=CC(C4=CC5=CC=CC=C5N=C4)=CC=C3N=C1)C(N2C6=CC=C(C=C6)C(C)(C#N)C)=O.CC7=CC=C(S(=O)(O)=O)C=C7 |
| Defactinib (hydrochloride) | -2.9 | 60.9 | 27.9 | 47.7 | O=C(NC)C1=CC=C(NC2=NC=C(C(F)(F)F)C(NCC3=NC=CN=C3N(C)S(=O)(C)=O)=N2)C=C1.[H]Cl |
| DGY-06-116 | 5.6 | 57.3 | 21.2 | 31.3 | O=C(NC1=C(Cl)C=CC=C1C)C(C=N2)=C(NC3=C(NC(C=C)=O)C=CC=C3)N=C2NC4=CC=C(N5CCN(C)CC5)C=C4 |
| Dinaciclib | 95.4 | 95.9 | 82.4 | 92.8 | OCC[C@H]1N(CCCC1)C2=NC3=C(C=NN3C(NCC4=C[N+]([O-])=CC=C4)=C2)CC |
| Doxorubicin (hydrochloride) | 58.9 | 46.5 | 1.7 | 41.1 | COC1=C2C(C(C(C(O)=C(C[C@](C(CO)=O)(O)C[C@@H]3O[C@@]4([H])C[C@H](N)[C@H](O)[C@H](C)O4)C3=C5O)=C5C2=O)=O)=CC=C1.[H]Cl |
| DS21360717 | -21.4 | 8.6 | 9.2 | 56.3 | N[C@H]1CCCC[C@H]1NC2=NC(NC3=CC(C)=CC=C3)=C4C(C=NNC4=O)=C2C#N |
| DTP3 (TFA) | 6.6 | 58.5 | 13.3 | 29.6 | O=C(N[C@H](CC1=CC=CC=C1)C(N)=O)[C@H](NC([C@H](NC(C)=O)CC2=CC=C(O)C=C2)=O)CCCNC(N)=N.OC(C(F)(F)F)=O |
| Dubermatinib | 42.4 | 42.3 | 58.0 | 41.2 | O=S(C1=CC=CC=C1NC2=NC(NC3=CC=C(CN4CCN(C)CC4)C=C3)=NC=C2Cl)(N(C)C)=O |
| eCF506 | 21.4 | 35.1 | 21.1 | 50.1 | NC1=C2C(C3=CC(OC)=C(NC(OC(C)(C)C)=O)C=C3)=NN(CCN4CCC(N(C)C)CC4)C2=NC=N1 |
| EGFR-IN-11 | 8.9 | 21.7 | 54.6 | 45.6 | CN1CCN(CC1)C2=CC=C(C=C2)NC3=NC4=C(C=N3)N=C(NC5=CC=CC=C5)N4[C@H]6CN(CC6)S(C7CC7)(=O)=O |
| Elimusertib | 35.9 | 24.4 | 42.1 | 50.9 | CN1N=CC=C1C2=C(C=CN=C3C4=CC=NN4)C3=NC(N5[C@H](C)COCC5)=C2 |
| Elimusertib (hydrochloride) | 33.1 | 34.4 | 50.0 | 51.1 | CN1N=CC=C1C2=C(C=CN=C3C4=CC=NN4)C3=NC(N5[C@H](C)COCC5)=C2.[H]Cl |
| ENMD-2076 | -9.9 | 50.2 | -24.1 | 8.2 | CC1=CC(NC2=NC(/C=C/C3=CC=CC=C3)=NC(N4CCN(CC4)C)=C2)=NN1 |
| ENMD-2076 (Tartrate) | 17.9 | 27.5 | 21.2 | 55.5 | O=C([C@@H]([C@H](C(O)=O)O)O)O.CC1=CC(NC2=NC(/C=C/C3=CC=CC=C3)=NC(N4CCN(CC4)C)=C2)=NN1 |
| ERK1/2 inhibitor 1 | 1.0 | 35.6 | -8.7 | 52.4 | ClC1=CN=C(NC2CCOCC2)N=C1C3=CC=C(CN([C@H](C)C(N[C@H](CO)C4=CC=CC(C)=C4)=O)C5=O)C5=C3 |
| ETP-46321 | 1.7 | 7.4 | 84.6 | 37.7 | NC(N=C1)=NC=C1C2=CN3C=C(CN4CCN(S(=O)(C)=O)CC4)N=C3C(N5CCOCC5)=N2 |
| Everolimus | 14.4 | 62.8 | 28.7 | 21.8 | O=C([C@@]1(O)[C@@H](CC[C@@H](C[C@@H](/C(C)=C/C=C/C=C/[C@H](C[C@@H](C)C([C@@H]([C@@H](/C(C)=C/[C@H]2C)O)OC)=O)C)OC)O1)C)C(N3CCCC[C@H]3C(O[C@@H](CC2=O)[C@@H](C[C@H]4C[C@H]([C@H](OCCO)CC4)OC)C)=O)=O |
| FAK inhibitor 2 | 36.7 | 62.4 | 22.3 | 60.9 | O=C(NC)C(C=CC=C1)=C1NC2=C(C(F)(F)F)C=NC(NC3=CC=C(C(N4CCN(C(SCCN(C)C)=S)CC4)=O)C=C3)=N2 |
| Fimepinostat | 71.1 | 69.6 | 36.2 | 73.4 | O=C(C1=CN=C(N(CC2=CC3=NC(C4=CC=C(OC)N=C4)=NC(N5CCOCC5)=C3S2)C)N=C1)NO |
| Flavopiridol (Hydrochloride) | 10.6 | 52.9 | 33.3 | 22.7 | O=C1C2=C(C=C(C([C@]3([H])[C@H](O)CN(C)CC3)=C2OC(C4=CC=CC=C4Cl)=C1)O)O.[H]Cl |
| FLLL32 | -19.2 | 30.1 | -10.2 | 60.8 | O=C(C1(C(/C=C/C2=CC=C(OC)C(OC)=C2)=O)CCCCC1)/C=C/C3=CC=C(OC)C(OC)=C3 |
| FMK | -0.7 | 54.0 | 8.2 | 27.0 | FCC(C1=C(C2=C(N=CN=C2N1CCCO)N)C3=CC=C(C=C3)C)=O |
| FRAX1036 | -0.3 | 54.4 | 12.9 | 30.1 | O=C1C(C2=CC=C(C3=NC(C)=CN=C3)C=C2Cl)=CC4=CN=C(NCCC5CCN(C)CC5)N=C4N1CC |
| GDC-0623 | 24.8 | 32.4 | 9.8 | 59.9 | FC(C=C(I)C=C1)=C1NC2=C(C(NOCCO)=O)C=CC3=CN=CN23 |
| GDC-4379 | 28.9 | 8.9 | 36.1 | 54.9 | O=C(C1=C2N=CC=CN2N=C1)NC3=CN(CC(N(C)C)=O)N=C3C4=CC(Cl)=CC=C4OC(F)F |
| Gliotoxin | -3.8 | 20.2 | 10.9 | 53.0 | O=C1N(C)[C@@]2(CO)C(N3[C@@]1(SS2)CC4=CC=C[C@H](O)[C@@]34[H])=O |
| GNF4877 | -23.2 | -3.5 | -14.8 | 66.5 | O=C([C@H]1CN(C2=C(NC(C3=NC(C4=CC(OC(C)C)=CC=C4F)=CN=C3N)=O)C=NC=C2)CCC1)O |
| GSK-1070916 | 35.6 | 51.3 | 5.5 | 28.6 | O=C(NC1=CC=C(C2=NN(C=C2C3=C4C(NC(C5=CC=CC(CN(C)C)=C5)=C4)=NC=C3)CC)C=C1)N(C)C |
| GSK2593074A | 5.3 | 17.1 | 24.1 | 56.6 | CN1N=CC(C2=CN=C(N)C3=C2SC=C3C4=CC5=C(N(C(CC6=CC=CC=C6)=O)CC5)C=C4)=C1 |
| GSK2606414 | 1.3 | 16.9 | 17.5 | 58.1 | NC1=C2C(N(C)C=C2C3=CC4=C(N(C(CC5=CC=CC(C(F)(F)F)=C5)=O)CC4)C=C3)=NC=N1 |
| GSK3182571 | 48.2 | 71.5 | 37.1 | 46.7 | O=C(NC)C1=CC=CC=C1NC2=NC(NC3=CC=C(N4CCN(CCCN)CC4)C=C3)=NC=C2Cl |
| GSK461364 | 69.1 | 94.7 | 71.6 | 85.5 | O=C(C(SC(N1C=NC2=CC=C(C=C12)CN3CCN(CC3)C)=C4)=C4O[C@@H](C5=CC=CC=C5C(F)(F)F)C)N |
| GSK-872 (hydrochloride) | -1.5 | 12.5 | 62.5 | 27.4 | O=S(C1=CC=C2N=CC=C(C2=C1)NC3=CC=C4SC=NC4=C3)(C(C)C)=O.[H]Cl |
| GW779439X | 96.3 | 97.4 | 45.5 | 89.5 | FC(C1=CC(NC2=NC=CC(C3=C4C=CC=NN4N=C3)=N2)=CC=C1N5CCN(C)CC5)(F)F |
| HDACs/mTOR Inhibitor 1 | 41.0 | 34.6 | 18.0 | 51.7 | O=C(NO)CCCCCCN1C2=NC(C3=CC=C(NC(N4C[C@@H](C)OCC4)=O)C=C3)=NC(N5CCOCC5)=C2C=N1 |
| Hesperadin | 52.9 | 39.9 | 24.4 | 49.6 | O=C1NC2=CC=C(C=C2/C1=C(NC3=CC=C(C=C3)CN4CCCCC4)\C5=CC=CC=C5)NS(CC)(=O)=O |
| Homoharringtonine | 84.0 | 95.1 | 65.3 | 80.1 | [H][C@@]12[C@](CCC3)(C=C(OC)[C@]2(OC([C@](CCCC(O)(C)C)(O)CC(OC)=O)=O)[H])N3CCC4=CC5=C(OCO5)C=C14 |
| Hydroxyfasudil | 5.3 | 58.2 | 14.2 | 21.1 | O=C1NC=CC2=C1C=CC=C2S(=O)(N3CCNCCC3)=O |
| Ifebemtinib | 4.0 | 14.6 | 22.3 | 57.7 | O=C(C1=C(F)C=C(NC2=NC=C(C(F)(F)F)C(OC3=CC=C4C=C3C(N(C4)C)=O)=N2)C(OC)=C1)NC5CCN(CC5)C |
| Ilginatinib | 10.4 | 18.3 | 72.0 | 36.7 | CN1N=CC(C2=CC(N[C@H](C3=CC=C(F)C=C3)C)=NC(NC4=NC=CN=C4)=C2)=C1 |
| iMDK (quarterhydrate) | 13.7 | 56.4 | 22.3 | 20.2 | O=C1C(C2=CN3C(SC(CC4=CC=C(F)C=C4)=C3)=N2)=CC5=CC=CC=C5O1.[1/4].O |
| JAK1-IN-8 | -6.7 | 60.6 | 15.8 | 36.9 | O=C(C1CC1)NC2=CN3C(C4=CC(F)=C(CN5CCS(=O)(CC5)=O)C=C4)=CC=CC3=N2 |
| K00546 | 48.5 | 38.0 | 72.0 | 52.5 | S=C(N1N=C(NC2=CC=C(S(=O)(N)=O)C=C2)N=C1N)NC3=C(F)C=CC=C3F |
| KG5 | 10.7 | 27.2 | -9.9 | 53.1 | CSC1=NC(OC2=CC=C(C=C2)C3=NN=C(N3)NC4=CC=CC(C(F)(F)F)=C4)=CC(N)=N1 |
| Lapatinib (ditosylate) | -4.3 | 42.3 | 7.7 | 60.4 | O=S(CCNCC1=CC=C(C2=CC=C3C(C(NC4=CC(Cl)=C(C=C4)OCC5=CC(F)=CC=C5)=NC=N3)=C2)O1)(C)=O.O=S(C6=CC=C(C)C=C6)(O)=O.O=S(C7=CC=C(C)C=C7)(O)=O |
| limertinib | 5.2 | 14.0 | 20.4 | 60.2 | O=C(NC1=CC(NC2=NC=C(C(NC3=CC4=C(C=CC=C4)C=C3)=N2)Cl)=C(C=C1N(CCN(C)C)C)OC)C=C |
| LM22A-4 | 7.0 | 50.0 | 18.8 | 3.9 | O=C(C1=CC(C(NCCO)=O)=CC(C(NCCO)=O)=C1)NCCO |
| Luxeptinib | 51.0 | 54.1 | 23.2 | 0.9 | O=C(NC1=C(F)C=C(F)C=C1F)NC2=CC(F)=C(C(C=CC(C3=NC(C)=CN3)=C45)=C5CNC4=O)C=C2 |
| MAX-40279 (hemiadipate) | 29.8 | 53.6 | 21.1 | 9.1 | O=C(O)CCCCC(O)=O.FC1=CC(OC)=C(C2=C(C)SC3=C2N=C(NC4=CN(C5CCNCC5)N=C4)N=C3)C=C1.[1/2] |
| Mevociclib | 83.4 | 96.3 | 66.8 | 82.9 | O=C(C1=NC=C(NC(/C=C/CN(C)C)=O)C=C1)N[C@@]2(C)CCC[C@@H](NC3=NC=C(Cl)C(C4=CNC5=C4C=CC=C5)=N3)C2 |
| Milciclib | 2.3 | 67.5 | 25.8 | 48.8 | O=C(NC)C1=NN(C2=C1C(C)(CC3=CN=C(N=C23)NC4=CC=C(C=C4)N5CCN(CC5)C)C)C |
| Mirdametinib | 33.9 | 47.8 | 30.7 | 66.3 | O=C(C1=CC=C(C(F)=C1NC2=CC=C(I)C=C2F)F)NOC[C@H](O)CO |
| Mitoxantrone | 62.4 | 68.3 | 60.4 | 67.7 | O=C1C2=C(C(NCCNCCO)=CC=C2NCCNCCO)C(C3=C(O)C=CC(O)=C13)=O |
| Mitoxantrone (dihydrochloride) | 55.8 | 54.0 | 57.8 | 61.7 | O=C1C2=C(C(NCCNCCO)=CC=C2NCCNCCO)C(C3=C(O)C=CC(O)=C13)=O.[H]Cl.[H]Cl |
| ML-099 | -10.8 | 24.3 | 55.1 | 32.7 | OC(C1=CC=CN=C1SCCC2=CC=CC=C2)=O |
| MLN0905 | 75.2 | 83.9 | 66.9 | 69.5 | CC1=NC=C(CCCN(C)C)C=C1NC2=NC(C(C=CC(C(F)(F)F)=C3)=C3NC(C4)=S)=C4C=N2 |
| Mps1-IN-3 (hydrochloride) | 15.4 | 63.1 | 33.6 | 40.6 | OC1CCN(C2=CC=C(NC3=NC(NC4=CC=CC=C4S(=O)(C(C)C)=O)=C5N=CNC5=N3)C(OC)=C2)CC1.[H]Cl |
| MPT0B392 | 62.9 | 58.0 | 27.1 | 20.2 | NC1=C2C(N=C(S(=O)(C3=CC(OC)=C(OC)C(OC)=C3)=O)C=C2)=CC=C1OC |
| MRTX1133 | 51.5 | 24.4 | -39.6 | 15.5 | OC1=CC(C2=C(F)C3=NC(OC[C@]45C[C@@H](F)CN4CCC5)=NC(N6C[C@H]7N[C@H](CC7)C6)=C3C=N2)=C8C(C#C)=C(F)C=CC8=C1 |
| Mutant EGFR inhibitor (NT-1) | 5.5 | 65.3 | 24.4 | 71.1 | C=CC(NC1=CC(NC2=NC=C(Cl)C(C3=CNC4=C3C=CC=C4)=N2)=C(OC)C=C1N(CCN(C)C)C)=O |
| Naquotinib (mesylate) | 2.6 | 64.9 | 23.9 | 53.7 | O=C(C1=NC(CC)=C(O[C@H]2CN(C(C=C)=O)CC2)N=C1NC3=CC=C(N4CCC(N5CCN(C)CC5)CC4)C=C3)N.CS(=O)(O)=O |
| Narciclasine | 67.8 | 80.0 | 59.8 | 74.8 | O=C1N[C@@]2([H])[C@H](O)[C@H](O)[C@@H](O)C=C2C3=C1C(O)=C4C(OCO4)=C3 |
| Neratinib | 3.1 | 34.4 | 9.7 | 57.4 | ClC1=C(OCC2=NC=CC=C2)C=CC(NC3=C(C#N)C=NC4=CC(OCC)=C(NC(/C=C/CN(C)C)=O)C=C43)=C1 |
| NVP-2 | 96.1 | 97.1 | 92.7 | 89.5 | ClC1=CN=C(N[C@H]2CC[C@H](N[C@H](C)COC)CC2)C=C1C3=CC=CC(NCC4(C#N)CCOCC4)=N3 |
| NVP-TAE 226 | -14.7 | 27.7 | -0.9 | 66.1 | O=C(C1=C(C=CC=C1)NC2=NC(NC3=C(C=C(C=C3)N4CCOCC4)OC)=NC=C2Cl)NC |
| Omipalisib | 45.2 | 61.7 | 17.2 | 78.9 | COC1=NC=C(C=C1NS(C2=CC=C(C=C2F)F)(=O)=O)C3=CC=C(C4=C3)N=CC=C4C5=CN=NC=C5 |
| ON-013100 | 10.1 | 59.0 | 18.3 | 29.1 | OC1=CC(CS(=O)(/C=C/C2=C(OC)C=C(OC)C=C2OC)=O)=CC=C1OC |
| Orantinib | -4.6 | 55.2 | 7.4 | 44.5 | CC1=C(/C=C2C3=CC=CC=C3NC/2=O)NC(C)=C1CCC(O)=O |
| OTS514 | 80.8 | 74.6 | 49.2 | 47.0 | O=C1NC2=C(C(C3=CC=C([C@@H](C)CN)C=C3)=C(O)C=C2C)C4=C1SC=C4 |
| OTSSP167 (hydrochloride) | 64.4 | 73.7 | 27.6 | 76.0 | ClC1=CC(C2=CC=C(N=CC(C(C)=O)=C3N[C@H]4CC[C@H](CN(C)C)CC4)C3=N2)=CC(Cl)=C1O.[H]Cl |
| Oxindole | 21.7 | -3.9 | 27.0 | 51.1 | O=C1NC2=C(C=CC=C2)C1 |
| p38? inhibitor 2 | -20.1 | 41.0 | -2.0 | 66.1 | O=C(NC1CC1)C2=CC=C(C)C(N3C(C(NC(C4=CC=CC=C4OCCNC)(C)C)=NC=C3)=O)=C2 |
| PD153035 | 4.8 | 63.8 | 12.3 | 1.5 | COC1=C(C=C(C2=C1)N=CN=C2NC3=CC(Br)=CC=C3)OC |
| PD318088 | 11.7 | 52.7 | 36.2 | 32.6 | BrC1=CC(C(NOCC(CO)O)=O)=C(C(F)=C1F)NC2=CC=C(C=C2F)I |
| PERK-IN-4 | -7.4 | 4.2 | 19.7 | 57.3 | NC1=C(C2=NC=N1)C(C3=CC=C4C(CCN4C(CC5=CC(C(F)(F)F)=CC(F)=C5)=O)=C3)=CN2C |
| PF-06409577 | 8.4 | 1.9 | 51.2 | 27.2 | O=C(C1=CNC2=C1C=C(C3=CC=C(C4(O)CCC4)C=C3)C(Cl)=C2)O |
| PF-6274484 | -6.1 | 7.7 | 23.1 | 50.6 | C=CC(NC1=CC2=C(NC3=CC(Cl)=C(F)C=C3)N=CN=C2C=C1OC)=O |
| PF-AKT400 | -4.9 | 26.9 | 30.5 | 66.7 | N[C@@](CC1)(CNC(C2=CC=C(F)C=C2F)=O)CN1C3=C4C(NC=C4CC)=NC=N3 |
| PHT-7.3 | 10.8 | 57.0 | 12.4 | 16.4 | O=C1C2=C(C=CC=C2)C(CC3OCCCO3)=NN1CC4=NC(C5=CC=C(C)C=C5)=CS4 |
| PI-828 | -12.6 | 29.9 | 57.4 | 12.5 | O=C1C=C(N2CCOCC2)OC3=C(C4=CC=C(N)C=C4)C=CC=C13 |
| Pictilisib | -0.1 | 58.9 | 28.1 | 15.8 | CS(N1CCN(CC2=CC3=C(C(N4CCOCC4)=NC(C5=CC=CC6=C5C=NN6)=N3)S2)CC1)(=O)=O |
| Pimasertib | 25.5 | 26.9 | 13.9 | 51.2 | FC1=C(C=CC(I)=C1)NC2=CN=CC=C2C(NC[C@@H](CO)O)=O |
| Prexasertib | 81.3 | 76.3 | 68.5 | 52.2 | NCCCOC(C=CC=C1OC)=C1C2=CC(NC3=NC=C(C#N)N=C3)=NN2 |
| Prexasertib (dihydrochloride) | 64.9 | 66.2 | 1.2 | 62.1 | NCCCOC(C=CC=C1OC)=C1C2=CC(NC3=NC=C(C#N)N=C3)=NN2.[H]Cl.[H]Cl |
| Prexasertib (dimesylate) | 66.1 | 68.2 | 64.8 | 62.8 | N#CC1=NC=C(NC2=NNC(C3=C(OC)C=CC=C3OCCCN)=C2)N=C1.CS(=O)(O)=O.CS(=O)(O)=O |
| Protein kinase inhibitor H-7 (dihydrochloride) | -6.1 | 53.0 | 17.4 | 27.5 | O=S(C1=CC=CC2=C1C=CN=C2)(N3C(C)CNCC3)=O.[H]Cl.[H]Cl |
| Quizartinib | -6.1 | 52.9 | 8.6 | 48.8 | O=C(NC1=NOC(C(C)(C)C)=C1)NC2=CC=C(C=C2)C3=CN4C(SC5=CC(OCCN6CCOCC6)=CC=C45)=N3 |
| RGB-286638 | 42.3 | 61.2 | 28.5 | 17.3 | O=C(NN1CCOCC1)NC2=CC=CC(C3=C4C(C5=CC=C(CN6CCN(CCOC)CC6)C=C5)=NN3)=C2C4=O.[H]Cl.[H]Cl |
| RIP2 kinase inhibitor 2 | -3.7 | 51.7 | -6.4 | 38.1 | CC(C)(C)S(C1=C(OCCOC)C=C(N=CC=C2NC3=NNC(C)=C3C)C2=C1)(=O)=O |
| Ro3280 | 33.2 | 55.1 | 18.9 | 37.6 | COC1=C(NC2=NC(N(C3CCCC3)CC(F)(F)C(N4C)=O)=C4C=N2)C=CC(C(NC5CCN(C)CC5)=O)=C1 |
| Roniciclib | 92.7 | 97.6 | 26.2 | 95.3 | C[C@@H](O)[C@H](OC1=NC(NC2=CC=C([S@@](=N)(C3CC3)=O)C=C2)=NC=C1C(F)(F)F)C |
| Sangivamycin | 63.3 | 94.3 | 37.7 | 27.2 | OC[C@@H]1[C@@H](O)[C@@H](O)[C@H](N2C3=NC=NC(N)=C3C(C(N)=O)=C2)O1 |
| Sapanisertib | 10.8 | 21.1 | 16.4 | 60.6 | NC1=NC=NC2=C1C(C3=CC4=C(C=C3)OC(N)=N4)=NN2C(C)C |
| SCH-1473759 (hydrochloride) | 65.2 | 75.7 | 26.6 | 45.7 | [H]Cl.CC1=CN2C(C(NC3=CC(CN(C(C)(CO)C)CC)=NS3)=N1)=NC=C2C4=CNN=C4 |
| SK-3-91 | 11.2 | 68.3 | 30.0 | 27.2 | O=C(N1C2C(NC(CC2)=O)=O)C3=C(NCC(NCCOCCN4CCN(C5=CC=C(NC6=NC(NC7=C(S(C(C)C)(=O)=O)C=CC=C7)=C(Cl)C=N6)C=C5)CC4)=O)C=CC=C3C1=O |
| SM1-71 | 65.5 | 67.9 | 47.4 | 60.4 | C=CC(NC1=CC=CC=C1NC2=NC(NC3=CC=C(N4CCN(C)CC4)C=C3)=NC=C2Cl)=O |
| SM-276001 | 5.7 | 51.7 | 4.4 | -4.5 | NC1=C2C(N(C(N2)=O)CC3=CN=C(C=C3C)C)=NC(NCCCC)=N1 |
| SOS1-IN-11 | -9.5 | 51.0 | 27.3 | 44.4 | CC1=NN=C(C2=CC(N3CCOCC3)=NC=C21)N[C@@H](C4=CC=CC(C(F)(F)F)=C4C)C |
| Staurosporine | 84.7 | 89.9 | 92.2 | 92.1 | O=C(NC1)C2=C1C3=C(C4=C2C5=C(C=CC=C5)N4[C@H]6C[C@@H](NC)[C@@H](OC)[C@]7(C)O6)N7C8=CC=CC=C83 |
| SU5214 | 6.7 | 55.5 | 19.4 | 20.7 | O=C1NC2=C(C=CC=C2)/C1=C/C3=CC=CC=C3OC |
| SY-5609 | 32.3 | 32.6 | 49.1 | 54.2 | N#CC1=C(P(C)(C)=O)C2=C(C=C1)C(C3=NC(N[C@@H]4CNC(C)(C)CC4)=NC=C3C(F)(F)F)=CN2 |

| | | | | | |
|---|---|---|---|---|---|
| T-00127_HEV1 | -2.1 | 51.3 | 5.7 | 7.1 | CC1=NC2=C(C3=CC=C(OC)C(OC)=C3)C(C)=NN2C(NCCN4CCOCC4)=C1 |
| TAK-593 | 3.4 | 59.0 | 3.9 | 21.3 | O=C(C1=CC(C)=NN1C)NC2=CC(OC3=NN4C(C=C3)=NC(NC(C5CC5)=O)=C4)=CC=C2C |
| TAK-733 | 35.7 | 38.0 | 24.2 | 60.5 | O=C1C(C(NC2=CC=C(I)C=C2F)=C(F)C(N3C)=O)=C3N=CN1C[C@@H](O)CO |
| TAK-901 | -4.2 | 21.4 | 18.8 | 69.1 | O=C(NC1CCN(CC1)C)C2=C(C3=C(C4=CC(C)=CN=C4N3)C(C5=CC=CC(S(=O)(CC)=O)=C5)=C2)C |
| TAK-960 | 68.7 | 63.6 | 59.2 | 40.7 | O=C(NC1CCN(C)CC1)C2=CC(OC)=C(NC3=NC=C(N4C)C(N(C5CCCC5)CC(F)(F)C4=O)=N3)C=C2F |
| TAK-960 (dihydrochloride) | 46.3 | 48.2 | 51.1 | 37.5 | [H]Cl.O=C(NC1CCN(C)CC1)C2=CC(OC)=C(NC3=NC=C(N4C)C(N(C5CCCC5)CC(F)(F)C4=O)=N3)C=C2F.[H]Cl |
| Talmapimod | -14.7 | 43.3 | -5.6 | 61.1 | O=C(N(C)C)C(C1=CN(C)C2=C1C=C(C(N3[C@H](C)CN(CC4=CC=C(F)C=C4)[C@@H](C)C3)=O)C(Cl)=C2)=O |
| Taselisib | 21.8 | 55.2 | 44.2 | 45.6 | CC(C)N1C(C2=CN3C(C(C(OCC3)=C4)=CC=C4C5=CN(C(C)(C(N)=O)C)N=C5)=N2)=NC(C)=N1 |
| Tauroursodeoxycholate | -30.8 | 45.2 | -1.2 | 51.7 | C[C@H](CCC(NCCS(=O)(O)=O)=O)[C@H]1CC[C@@]2([H])[C@]3([H])[C@@H](O)C[C@]4([H])C[C@H](O)CC[C@]4(C)[C@@]3([H])CC[C@]12C |
| Tirbanibulin | 36.9 | 38.9 | 11.7 | 56.7 | O=C(CC1=NC=C(C2=CC=C(OCCN3CCOCC3)C=C2)C=C1)NCC4=CC=CC=C4 |
| Tirbanibulin (Mesylate) | 41.2 | 32.4 | 21.9 | 58.3 | O=C(NCC1=CC=CC=C1)CC2=NC=C(C3=CC=C(OCCN4CCOCC4)C=C3)C=C2.CS(=O)(O)=O |
| Torin 2 | 53.7 | 61.4 | 18.9 | 52.7 | O=C1N(C2=C(C=C1)C=NC3=CC=C(C=C32)C4=CC=C(N=C4)N)C5=CC=CC(C(F)(F)F)=C5 |
| Toyocamycin | 81.3 | 91.8 | 77.6 | 91.6 | O[C@H]1[C@@H](O[C@H](CO)[C@H]1O)N2C3=C(C(N)=NC=N3)C(C#N)=C2 |
| TPCA-1 | -20.4 | -0.9 | 56.7 | 29.4 | O=C(C1=C(NC(N)=O)SC(C2=CC=C(F)C=C2)=C1)N |
| Trametinib (DMSO solvate) | 36.1 | 36.6 | 49.4 | 55.1 | CN1C(C(C)=C(N(C(N(C2=O)C3CC3)=O)C4=CC=CC(NC(C)=O)=C4)C2=C1NC5=CC=C(C=C5F)I)=O.CS(C)=O |
| Vevorisertib (trihydrochloride) | 6.4 | 63.8 | 16.2 | 36.3 | NC1=C(C=CC=N1)C2=NC3=CC=C(C4=CC(N(CC5)CCC5N(C)C(C)=O)=CC=C4)N=C3N2C6=CC=C(C7(CCC7)N)C=C6.[H]Cl.[H]Cl.[H]Cl |
| Volasertib | 43.6 | 60.1 | 33.8 | 15.5 | O=C(C1=CC=C(C(OC)=C1)NC2=NC=C(N(C3=O)C)C(N([C@@H]3CC)C(C)C)=N2)N[C@H]4CC[C@@H](CC4)N5CCN(CC5)CC6CC6 |
| WNK-IN-11 | -31.2 | 35.1 | -38.5 | 50.8 | O=C(C1=CC(C2=CSC(NC)=N2)=NC=C1Cl)N3CCN(CC4=CC=C(Cl)C=C4)CC3 |
| XRP44X | 64.1 | 70.4 | 55.9 | 62.9 | O=C(N1CCN(C2=CC(Cl)=CC=C2)CC1)C3=CC(C)=NN3C4=CC=CC=C4 |
| YKL-06-062 | 9.6 | 56.9 | 19.1 | 66.5 | O=C1N(C2CCCCC2)C3=NC(NC4=CC=C(N5CCN(C)CC5)C=C4)=NC=C3CN1C6=C(C)C=CC=C6C |
| YKL-5-124 | 41.5 | 72.9 | 30.8 | 47.1 | CC1(C)N(C(N[C@@H](C2=CC=CC=C2)CN(C)C)=O)CC3=C1NN=C3NC(C4=CC=C(NC(C=C)=O)C=C4)=O |
| YKL-5-124 (TFA) | 36.9 | 58.8 | 32.3 | 39.6 | CC1(C)N(C(N[C@@H](C2=CC=CC=C2)CN(C)C)=O)CC3=C1NN=C3NC(C4=CC=C(NC(C=C)=O)C=C4)=O.FC(F)(C(O)=O)F |
| Zanubrutinib | -11.6 | 56.7 | 8.4 | 16.4 | NC(C1=C2N(N=C1C3=CC=C(OC4=CC=CC=C4)C=C3)[C@H](C5CCN(C(C=C)=O)CC5)CCN2)=O |
| ZD-4190 | -3.4 | 14.5 | 71.1 | 37.7 | COC1=CC2=C(NC3=CC=C(Br)C=C3F)N=CN=C2C=C1OCCN4N=NC=C4 |
| ZM-447439 | -1.5 | 51.8 | 11.1 | 47.5 | O=C(NC1=CC=C(NC2=C(C(C=C3OCCCN4CCOCC4)=NC=N2)C=C3OC)C=C1)C5=CC=CC=C5 |
| ZT-12-037-01 | -1.0 | 18.0 | 11.6 | 74.6 | COC1=CC2=NC(NC3CC3)=NC(NC4CCN(C(C)C)CC4)=C2C=C1OC |

**Y: Hit**, N: not hit

| Cat. (#) | Monotherapy | | Combination | | Interpretation |
|---|---|---|---|---|---|
| | Parental | Resistant | Parental | Resistant | |
| **1** | Y | Y | Y | Y | General cytotoxicity |
| **2** | N | N | Y | Y | General KRASi **synergizer** |
| **3** | N | N | N | Y | Resistant cell selective KRASi **synergizer** |
| **4** | N | N | Y | N | Parental cell selective KRASi **synergizer** |
| **5** | Y | N | Y | Y | Parental & Resistant cell selective KRASi **synergizer** |
| **6** | N | Y | N | Y | KRASi resistant cell selective hits |
| **7** | N | Y | Y | Y | KRASi resistant cell selective hits |
| **8** | Y | N | Y | N | Parental selective, KRASi resistance dependent resistance |
| **9** | N | Y | Y | N | Unlikely |
| **10** | N | Y | N | N | Unlikely |
| **11** | Y | Y | N | N | Unlikely |
| **12** | Y | N | N | N | Unlikely |
| **13** | Y | N | N | Y | Unlikely |
| **14** | Y | Y | N | Y | Unlikely |
| **15** | Y | Y | Y | N | Unlikely |
| **16** | N | N | N | N | Not hit |
